# Calcium imaging in freely-moving mice during electrical stimulation of deep brain structures

**DOI:** 10.1101/460220

**Authors:** J. K. Trevathan, A. J. Asp, E. N. Nicolai, J. M. Trevathan, N.A. Kremer, T.D. Kozai, D. Cheng, M. Schachter, J. J. Nassi, S. L. Otte, J. G. Parker, J. L. Lujan, K. A. Ludwig

## Abstract

After decades of study in humans and animal models, there remains a lack of consensus regarding how the action of electrical stimulation on neuronal and non-neuronal elements – e.g. neuropil, cell bodies, glial cells, etc. – leads to the therapeutic effects of neuromodulation therapies. To further our understanding of neuromodulation therapies, there is a critical need for novel methodological approaches using state-of-the-art neuroscience tools to study neuromodulation therapy in preclinical models of disease. In this manuscript we outline one such approach combining chronic behaving single-photon microendoscope recordings in a pathological mouse model with electrical stimulation of a common deep brain stimulation (DBS) target. We describe in detail the steps necessary to realize this approach, as well as discuss key considerations for extending this experimental paradigm to other DBS targets for different therapeutic indications. Additionally, we make recommendations from our experience on implementing and validating the required combination of procedures that includes: the induction of a pathological model (6-OHDA model of Parkinson’s disease) through an injection procedure, the injection of the viral vector to induce GCaMP expression, the implantation of the GRIN lens and stimulation electrode, and the installation of a baseplate for mounting the microendoscope. We proactively identify unique data analysis confounds occurring due to the combination of electrical stimulation and optical recordings and outline an approach to address these confounds. In order to validate the technical feasibility of this unique combination of experimental methods, we present data to demonstrate that 1) despite the complex multifaceted surgical procedures, chronic optical recordings of hundreds of cells combined with stimulation is achievable over week long periods 2) this approach enables measurement of differences in DBS evoked neural activity between anesthetized and awake conditions and 3) this combination of techniques can be used to measure electrical stimulation induced changes in neural activity during behavior in a pathological mouse model. These findings are presented to underscore the feasibility and potential utility of minimally constrained optical recordings to elucidate the mechanisms of DBS therapies in animal models of disease.

## Introduction

A multitude of neurologic and psychiatric disorders affect up to one billion people worldwide^1^, and up to thirty percent of the patients diagnosed with neurologic disorders remain refractory to conventional pharmacologic, behavioral, and surgical interventions^2,3^. To address this patient population, the last 30 years have seen a significant expansion in the application of devices to electrically stimulate the nervous system for therapeutic effect, known as neuromodulation therapies, that have become established forms of treatment for patients suffering from these neurologic and psychiatric conditions. For example, electrical stimulation of deep brain structures, known as deep brain stimulation (DBS), has received approval from the United States Food and Drug Administration (FDA) for the treatment of advanced Parkinson’s disease (PD), tremor^4–6^, and epilepsy^7–10^. Additionally, humanitarian device exemptions have been granted for treatment of dystonia and obsessive-compulsive disorder^6^. Despite the clinical prevalence of DBS therapies, the mechanisms of action for any specific therapeutic use of DBS are poorly understood.

Studies to elucidate mechanisms of action of DBS have centered on measuring the evoked physiological responses using non-invasive functional imaging^11–14^ and electrophysiological recording techniques^15–20^. Unfortunately, these techniques have inherent limitations that make it difficult to parse out the multimodal mechanisms by which DBS may alter the activity of a neural circuit during the execution of complex behavior. Functional imaging techniques - such as functional magnetic resonance imaging (fMRI) and positron emitted tomography (PET) - have low spatial (on the order of a millimeter) and temporal resolution (on the order of several seconds) and can only be performed during simple behavioral paradigms that restrict overall movement^21,22^. Electrophysiological recordings from single-units or multi-unit populations provide high spatial and temporal resolution but are limited to sparse sampling (typically under 100 electrode contacts separated by hundreds of microns), provide limited or unreliable information about recorded cell-types and their exact spatial locations^23^, and have low signal stability over time that limits the study of DBS induced plastic changes in circuit function^24,25^. As DBS induces electromagnetic cross-talk and disturbs the recording electrode/electrolyte interface, electrical stimulation also generates spurious artifacts that can resemble electrophysiological responses^26,27^.

The intended therapeutic and unintended side effects of DBS are likely mediated through multiple mechanisms, potentially driven by multiple cell-types, that alter the function of a neural circuit. The relative contribution of each mechanism may evolve over the course of stimulation due to changes in the electrode/tissue interface^28,29^, stimulation induced plasticity in neuronal and non-neuronal cells near the electrode^30^, stimulation induced post-synaptic plasticity^31^, changes in behavior state^32^, and adjunct drug status^33^. Clinical use of neuromodulation therapies in diverse conditions such as epilepsy, depression and Tourette’s syndrome consistently demonstrate significant improvement in patient outcomes with slow onsets over periods of several weeks to several months^11,34–42^. Whether these slow therapeutic responses are a result of placebo effect, changes to tissue local to the implant, wider network adaptation, or combinations of all the above, remains unknown. Consequently, experimental paradigms for elucidating the therapeutic mechanisms of DBS ideally need to be able to record from stable populations of hundreds of genetically specified cells with high spatial and temporal resolution across timeframes of weeks to months. These measurements need to be made during behaviors relevant to the clinical indication being studied to understand the role of both the local, circuit, and systems level dependencies in the neural response to DBS.

Single-photon calcium imaging via a head-mounted miniature microscope has previously been demonstrated in behaving animals to record from large stable populations of genetically specified cells across timeframes of weeks to months^43–46^ but has yet to be extended to the study of neuromodulation therapies. Towards that end, here we introduce our method for combining electrical stimulation of deep brain structures with chronic single-photon imaging in a behaving mouse with induced disease pathology (Figure 1A,B). Combining single-photon microscopy in a pathological model of DBS adds several invasive components to the experimental procedure, which introduce more potential failure points. Consequently, we provide recommendations from our experience in implementing this combination of methods on how to separately refine and validate each component step to achieve a combined procedure. As synchronous neural activity evoked by electrical stimulation confounded some state-of-the-art miniature microscopy analysis techniques, we present a qualitative assessment of different analysis techniques to underscore the importance validating the accurate identification of neuronal calcium traces in the context of neuromodulation. As the integration of these multiple complex components may impact recording quality, we provide data to verify the technical feasibility of maintaining stable optical recordings of hundreds of striatal cells for week long periods in the 6-OHDA mouse model of Parkinson’s Disease also implanted with electrodes to provide stimulation to the subthalamic nucleus (Figure 1C). We also present our initial within-animal findings that STN stimulation evoked responses significantly differ as a function of anesthetized versus awake states, as well as between rest versus movement in the awake state. These initial data illustrate the need for studying DBS mechanisms in awake, behaving pathological models. Whether these within-animal differences elucidate fundamental mechanisms of DBS therapies across a cohort will be explored in future studies.

**Figure 1.**
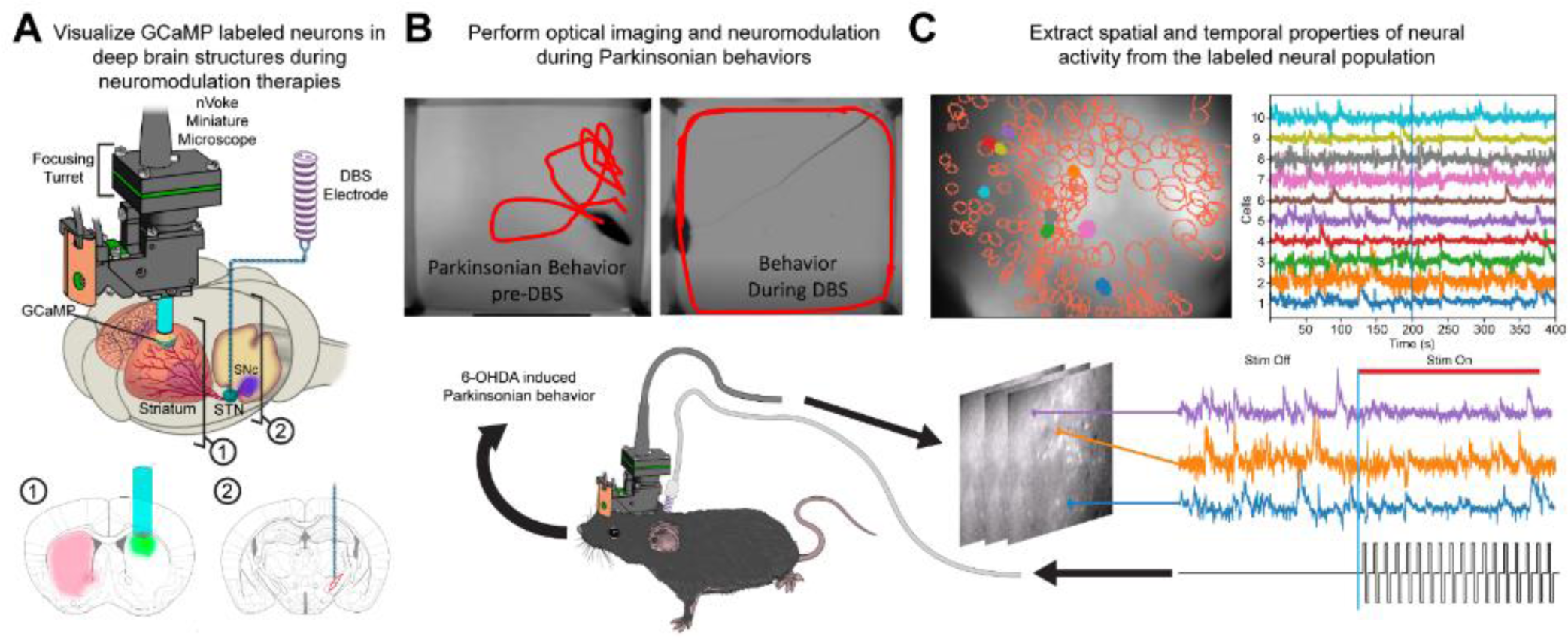
Experimental objectives. **A)** Visualization of GCaMP labeled neurons in deep brain structures of 6-hydroxy dopamine (6-OHDA) lesioned mice was performed with a miniature head-mounted microscope (Inscopix, Inc., Palo Alto, CA). The substantia nigra pars compacta (SNc) was lesioned via an injection of 6-hydroxydopamine (6-OHDA) to create an animal model of Parkinson’s disease (PD). Calcium imaging of cells labeled with GCaMP, a genetically encoded calcium indicator commonly used to detect neural activity, was performed through a gradient index (GRIN) lens implanted into the striatum (1). Additionally, a twisted bipolar electrode was inserted into the subthalamic nucleus (STN) to deliver deep brain stimulation (DBS) (2). **B)** Optical imaging and electrical stimulation of the STN were performed during open field behavior while position of the animal was tracked (red lines) to assess behavioral changes. **C)** Spatial (outlines) and temporal (traces) properties of neural activity, shown on top left and right panels, respectively, were extracted from the recorded miniature microscopy videos and analyzed using constrained non-negative matrix factorization for microendoscope data (CNMF-E) to determine changes in striatal activity that occurred during stimulation (bottom panel). This figure is not covered by the CC-BY license. Used with permission of Mayo Foundation for Medical Education and Research. All rights reserved.

## Methods

The experimental protocol to obtain calcium imaging in a behaving 6-OHDA lesioned mouse with electrical stimulation required the successful implementation of four separate components: 1) induction of calcium indicator expression via a unilateral injection of an AAV9 viral vector into striatum (Figure 2A,B), 2) placement of a Gradient Refractive Index (GRIN) lens into the target recording location just above striatum (Figure 2A,C), 3) induction of a Parkinsonian phenotype via unilateral injection of 6-OHDA into the substantia nigra pars compacta (SNc) (Figure 2A,B), and 4) placement of a bipolar stimulating electrode within the STN for application of electrical stimulation (Figure 2A,B). Each component of this experimental protocol was first tested in isolation in order to minimize the time and troubleshooting needed to implement the complete experiment.

**Figure 2.**
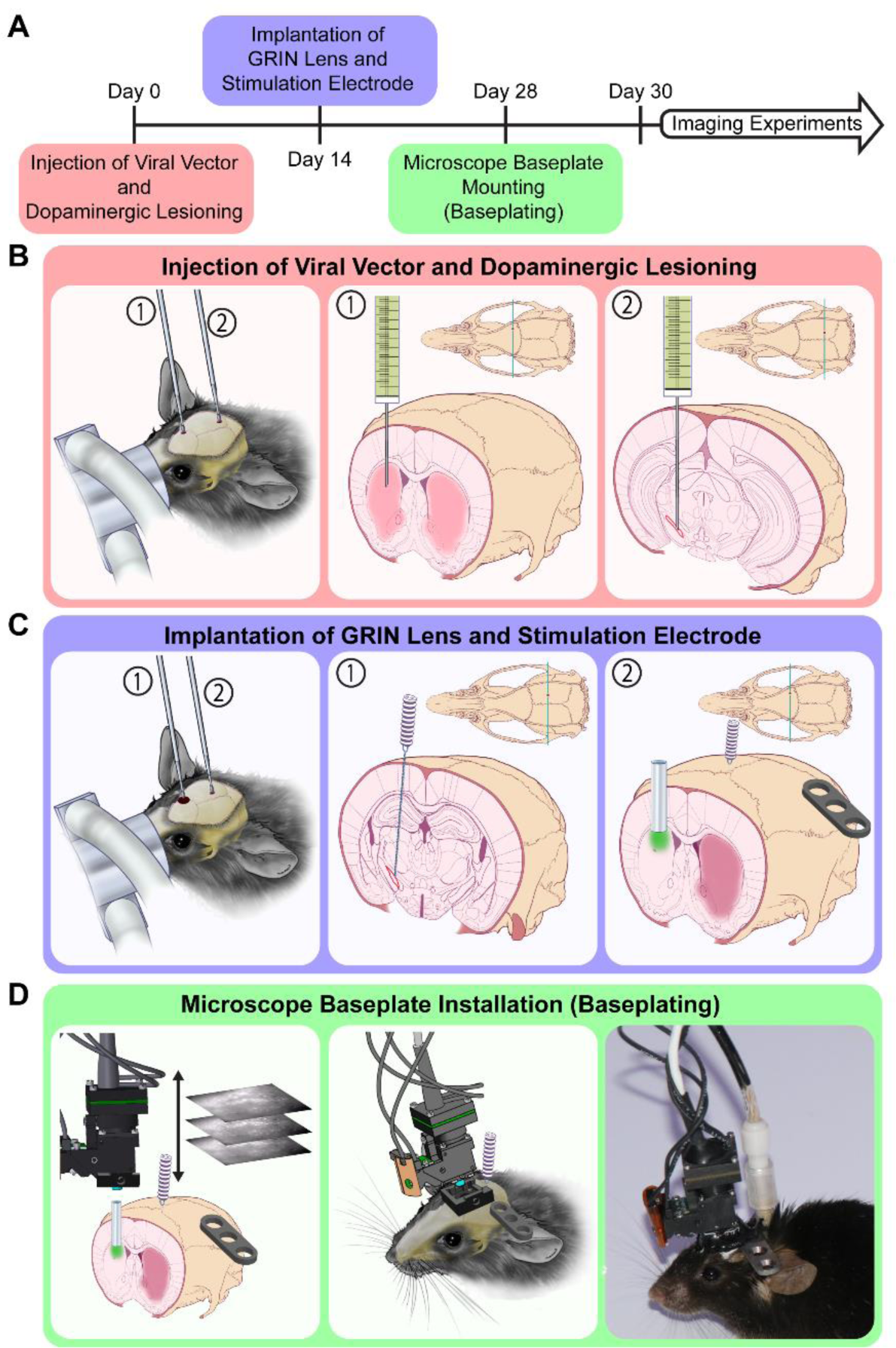
Experimental procedures required for calcium imaging of striatal neural activity during electrical stimulation of the subthalamic nucleus (STN) in awake, freely-behaving Parkinsonian animals. **A)** Mice underwent staged dopaminergic lesioning, viral transfection, surgical implantation of Graded Index (GRIN) lens and deep brain stimulation (DBS) electrode, and mounting of the microscope baseplate procedures over the course of one month prior to the start of calcium imaging experiments. **B)** At day 0, a unilateral microinjection of AAV9-GCaMP6m was performed into the dorsal striatum under isoflurane anesthesia to induce expression of the GCaMP reporter needed for imaging (1). In the same surgical session, a solution of 6-OHDA was injected ipsilaterally into the substantia nigra pars compacta (SNc) to cause a dopaminergic lesion, thereby resulting in a parkinsonian phenotype (2). **C)** At day 14, a DBS electrode (1) and GRIN lens (2) were implanted into the STN and dorsal striatum, respectively under isofluorane anesthesia. Both implants were secured to the skull via skull screws and dental adhesive. Extra care was placed to avoid covering bregma and the GRIN lens location. The panel on the right shows the baseplate used to mount the microscope. **D)** The microscope, with a baseplate attached, was positioned above the GRIN lens of an awake head-fixed animal positioned on a running wheel (not shown) to encourage movement. The distance from the microscope to GRIN lens was adjusted to optimize the focal plane within the tissue that exhibited the most active and in-focus cell bodies. The microscope baseplate was then affixed to the animal via dental cement and the headcap was coated with black nail polish to prevent contamination from external light sources. The animal was then removed from the head-fixed running wheel and ready for behaving imaging sessions. This figure is not covered by the CC-BY license. Used with permission of Mayo Foundation for Medical Education and Research. All rights reserved.

### Subjects

In total 30 adult C57BL/6J male mice approximately 25-30g (Jackson Laboratories, Bar Harbor, ME) were used to test each component of the experimental protocol. These animals were used for verifying 6-OHDA lesioning and behavior (n=15), and chronic recordings in healthy mice and parkinsonian mice (n = 5 and n=10, respectively). Of the ten parkinsonian mice, four were removed from analysis due to surgical complications and headcap failure while the others were used for data collection while under anesthesia (n=6). Two these six animals in which anesthetized recordings were performed were removed from the analyses presented here, as post-mortem histology indicated their electrodes were not properly placed within STN. Awake calcium imaging studies were also performed in two of these animals.

Animals were group-housed and maintained under a standard light/dark cycle with ad libitum access to food and water. All experimental procedures were approved by the either the Mayo Clinic Institutional Animal Care and Use Committee (IACUC) or the IACUC at LifeSource Biomedical Services, NASA Ames Research Center, California and were carried out in accordance with the U.S. National Research Council Guide for the Care and Use of Laboratory Animals.

### Injection of Viral Vector and Dopaminergic Lesioning

Mice were anesthetized in an induction chamber with 5% isoflurane (inhalation) and then maintained with a continuous flow of 1-2% isoflurane-oxygen mixture during stereotactic surgery (Kopf Systems, Tujunga, CA). A midline incision was performed to expose the skull and burr holes were made above the dorsal striatum (ML=1.65, AP=1.0 from bregma) and substantia nigra pars compacta (SNc) (ML=1.25, AP=–3.5 from bregma). A Hamilton syringe (Hamilton Company, Reno, NV) controlled by a stereotactically mounted microdrive (WorldPrecision Instruments) was lowered into right dorsal striatum (AP= +1.0 mm, ML=+1.65 mm, DV= −2.7 mm from dura) at an approximate rate of 10µm/sec and used to inject 500 nL of AAV9.CAG.GCaMP6m.WPRE.bGHpA (titer = 5.5E12 GC/mL; SignaGen Laboratories, Rockville, MD) at a rate of 100 nL/min (Figure 2B). Brain tissue was given one minute to settle prior to the unilateral AAV injection. The virus was allowed to diffuse for an additional five minutes prior to removing the needle at approximately 10µm/sec.

Immediately following AAV injection, a separate Hamilton syringe was used to inject 6-OHDA (4 µg/µL) into the SNc at three depths: DV –4.2, −4, and −3.8 from dura (Figure 2B). Injections at each location were 650 nL, and delivered at a rate of 100 nL/min. At each of the three depths, the injection was preceded with a wait period of one minute to allow brain tissue to stabilize, and followed by a wait period of five minutes for diffusion^43^. Insertion, removal and movements of the injection needle were carried out at approximately 10µm/sec. Desipramine HCl was injected (25 mg/kg, i.p.) approximately 30 minutes before the first 6-OHDA injection to prevent uptake of 6-OHDA into monoamine neurons^47^. The burr holes were sealed with Kwik-Sil (World Precision Instruments, Sarasota, FL) before suturing the skin incision. Ketofen (2.5mg/kg, i.p.) and Carprofen (2.5mg/kg, i.p.) were administered to reduce inflammation and provide analgesia. Each animal was allowed to recover individually in a heated home cage until fully mobile and were then returned to social housing.

### Implantation of GRIN Lens and DBS Electrode

Since the electrodes used for application of electrical stimulation to the STN and the GRIN lens needed to be embedded in the animal’s dental cement headcap, both were inserted during the same surgical procedure. Detailed procedures for implantation of the GRIN lens, on its own, are well described in Resendez et al. 2016^48^ and the GRIN lens implant location for striatal imaging was selected in accordance with Parker et al 2018^43^. Although DBS electrode implant sites, electrode sizes and stimulation parameters have been described in rat models^49^, much less work has been done in mice. For this reason, STN stimulation electrode sizes and the implant location were adapted from relevant rat studies^50,51^ based on the mouse brain atlas^52^. To ensure adequate time for viral transfection,^43^ two weeks after injection of viral vector and dopaminergic lesioning animals were prepared for a second surgery as described above. Burr holes were drilled above the dorsal striatum (ML 1.65, AP 1.0 from bregma) and STN (ML 1.45, AP –2.0 from bregma) (Figure 2C). Three bone screws (Component Supply Company, Sparta TN) were placed to add structural support for the C&B Metabond (Parkell, Edgewood, NY) dental cement used to secure the implants. The dura was surgically opened through the STN burr hole using microdissection forceps to allow insertion of the electrode. A twisted bipolar teflon-insulated platinum DBS electrode (Plastics One, Roanoke, VA) with a 500 µm exposed tip, and contacts separated to 500 µm apart, was inserted to the STN (ML 1.45, AP –2.0, DV −4.7 from dura) at approximately 10 µm/sec (Figure 2C). Through the striatum burr hole, a 1 mm diameter column of tissue above the dorsal surface of the striatum was then aspirated to clear a path for lens implantation. To prevent aspirating deeper than the intended target, a blunt 27-gauge needle was bent 2.1 mm from the tip, which is approximately the depth from the surface of the skull to the top of the dorsal striatum, and used to aspirate tissue until the white matter of the corpus collosum was visually identified through a Leica M320 surgical microscope (LEICA, Wetzlar, Germany). A small portion of white matter was then carefully removed through aspiration to expose the dorsal surface of the striatum for imaging while minimizing tissue damage. A 4 mm long, 1 mm diameter GRIN lens (Inscopix, Palo Alto, CA) was then inserted through the striatal burr hole to a depth of - 2.35 mm from dura using a custom-built lens holder (Figure 2C). The custom-built lens holder was fabricated from a micropipette tip cut so that ~500µm of the lens was securely held. Kwik-Sil was used to seal both burr holes following implantation of the GRIN lens and DBS electrode. A headbar was mounted so that it extended laterally from the animal’s skull contralateral to the implants to allow head-fixing of the animal while installing the microscope baseplate (Figure 2D). C&B Metabond was used to secure the headbar, DBS electrode, and GRIN lens to the skull forming a headcap. The lens surface was protected using a custom-made plastic cap made from the bottom tip of a microcentrifuge tube attached with Kwik-Cast (World Precision Instruments, Sarasota, FL). The same post-operative approach from the previous section was followed.

### Microscope Baseplate Installation (Baseplating)

In order to perform calcium imaging during behavior with the nVoke miniature microscope (Inscopix, Palo Alto, CA), a mounting baseplate attached above the GRIN lens is needed^48^. Baseplates were permanently fixed to the headcap of the mouse at the position that maximized the number of active in focus neurons. Due to the locomotion dependence of striatal neural activity^43,53,54^, the position was determined by visualizing active neurons in awake head-fixed mice. This procedure was completed two weeks after the GRIN lens implantation to allow for recovery (Figure 2A). Mice were first head-fixed using the previously attached headbar and positioned on a custom-built running wheel to allow locomotion. Locomotion increases striatal neuron activity, making it easier to determine the mounting position that maximizes the number of in focus neurons^43,53^. The microscope focusing turret (Figure 1A) was initially set near the middle of its range to maximize the ability to correct for movement of the baseplate in either direction that can occur during dental cement curing. Next, with a baseplate attached, the nVoke microscope was positioned above the surface of the lens (Figure 2D). The microscope was then leveled relative to the surface of the GRIN lens by ensuring the entire circumference around the edge of the lens was in focus. The excitation LED was then turned on to allow visualization of the fluorescent indicator^48^. The dorsoventral position of the miniature microscope was slowly varied over the area of tissue with the largest calcium signal until an imaging plane with in-focus active cells was found (Figure 2D). The mediolateral and anteroposterior positions of the microscope were adjusted as necessary to maximize the number of active cells in the field of view. Once this position was found, the baseplate was attached to the headcap using C&B Metabond. Finally, the headcap was painted with black nail polish to prevent ambient light from filtering through the dental cement and contaminating the recordings (Figure 2D).

### Calcium Imaging in Anesthetized Mice

Isoflurane was used to perform calcium imaging in anesthetized mice. The microscope was then attached to the baseplate, and the position of the microscope turret was adjusted to the imaging plane found during the baseplating procedure. This new turret position was recorded and used for all future imaging sessions. A series of 10-second duration, 130 Hz biphasic stimulation trains with 140 us pulse width were applied in one-minute intervals using a Master-8 stimulator with Iso-Flex isolators (A.M.P.I, Jerusalem, Israel). The amplitude of the stimuli was increased until evoked changes in calcium signal were observed. Stimulation trains were then applied at frequencies of 30 Hz, 80 Hz, and 130 Hz and repeated 3 times^32^. These stimulation frequencies were selected based on previous studies in 6-OHDA lesioned rat models showing therapeutic effects at high frequency (80 Hz and 130 Hz) but not low frequency (30Hz) stimulation^32^. Stimulation was triggered and synchronized with optical recordings using the digital I/O from the nVoke microscope system.

### Open Field Calcium Imaging During DBS

Mice were placed in a 45 cm by 45 cm open field arena (Med Associates, Fairfax, VT) with the miniature microscope and DBS electrodes connected. Animal movement was recorded with an overhead camera. The nVoke microscope sync signal provided through the nVoke digital IO box was used trigger to frame acquisition on the overhead camera. In non-lesioned animals without DBS implants, a 30-minute recording was completed two days after mounting the microscope baseplate and was repeated weekly thereafter for four weeks. In 6-OHDA lesioned mice with DBS electrodes implanted, a DBS parameter selection session similar to the previously described session under anesthesia was completed four days after mounting the microscope baseplate. During the parameter selection session 1-minute stimulation trains 130 Hz biphasic pulses (140 µs) were applied at 3-minute intervals, while simulation amplitude was increased to determine the stimulation threshold for evoking adverse effects. The most commonly observed adverse effect was abnormal motor activity or muscle contraction. Two 50-minute recordings were then performed on days 6 and 8 after baseplating.

### Identification of Neural Signals

Calcium imaging data were loaded into Inscopix Data Processing Software (IDPS) (Inscopix, Palo Alto, CA) for preprocessing prior to identification of neural signals via constrained non-negative matrix factorization for microendoscope data (CNMF-E)^55,56^. All data were first spatially down sampled by a factor of two, which is common practice in order to decrease processing time for subsequent analyses^48,57^. It was not necessary to use motion correction algorithms on data sets taken from anesthetized animals. However, calcium imaging data sets recorded in awake animals were then motion corrected with the Turboreg motion correction algorithm^58^ as implemented in IDPS. To improve processing time and memory usage during the application of CNMF-E, data were downsampled within IDPS by an additional factor of three for a total effective downsampling of 6x. Data were then exported as TIF files from IDPS and loaded into Matlab® (Natick, MA). To identify neurons, the Matlab implementation of CNMF-E (https://github.com/zhoupc/CNMF_E) was applied to the data following the recommended procedure from Zhou et al. (2018)^55^ and computation was performed on the Neuroscience Gateway^59^ high performance computing resource. Briefly, analysis was performed in patch mode using the largest possible patch sizes based on the available memory (128 GB) of the computational node. The ring model of background fluorescence^55^ was used for all data sets. Neuron and ring sizes were selected based on the data set so that ring diameter was approximately 1.5 times the largest neuron diameters and described n Zhou et al. (2018)^55^.

### Evaluation of Calcium Event Rate

To demonstrate the utility of this combination of methods in the study of DBS in freely behaving mice, changes in the calcium event rate normalized to rest were calculated from data collected during open field behavior and analyzed using CNMF-E. The calcium event rate was calculated and normalized to the calcium event rate of the same animal at rest as described in Parker et al^43^. Briefly, the instantaneous calcium event rate for each frame was calculated as the total number of events, as output by CNMF-E, occurring in the frame. Frames were then binned based on locomotor velocity, as assessed using Ethovision XT (Noldus, Wageningen, The Netherlands). Calcium event rates were normalized to the mean event rate at rest to compare results between animals, which was defined as when the instantaneous velocity of the animal was less than 0.5 cm/s^53^. We then binned the calcium event rates into sets of values for which the locomotor velocity was the same to within 0.5–2.5 cm/s, using finer bins at lower velocity and coarser bins at higher velocity^43^. For calcium event rate calculations during the application of deep brain stimulation, the calcium event rates during ambulation (locomotor velocity > 0.5 cm/s) were normalized to the mean event rat in the same animal at rest during unstimulated periods of the recording.

### Perfusion and Tissue Preparation

Just prior to euthanasia with a lethal dose of Euthasol (Virbac AH, Inc., Fort Worth, TX), electrochemical impedance spectroscopy analysis of each stimulation electrode contact was performed using an Autolab PGSTAT302N (Metraohm Autolab, Utrecht, The Netherlands) and following the spectroscopy sweep described in Lempka et al. 2009^60^. Animals with stimulation electrodes exhibiting impedances greater than 120kΩ of impedance at 130Hz were removed from further analysis due to failure of the stimulation electrode. Following euthanasia, transcranial perfusion was performed with 50mL of 0.9% saline, followed by 50mL of 4% paraformaldehyde in 0.1M NaKPO_4_. Perfused brains were extracted, post-fixed overnight in 4 % paraformaldehyde, and stored in 30% glycerol sinking solution. Cryosectioning with 40 μm slices was performed on a sliding microtome (Leica Biosystems, Wetzlar, Germany). Sections were stored in 0.1 % Na Azide in 0.1 M phosphate buffer solution.

### Histological Evaluation of 6-OHDA Lesion and Stereotactic Targeting

Coronal slices were rinsed in 0.01 M Phosphate buffered saline (PBS) with 0.2% Triton X-100 (PBS-Tx) and blocked in 10% normal goat serum in PBS-Tx for one hour. Slices were then incubated with the primary antibody, anti-Tyrosine Hydroxylase (rabbit polyclonal, Abcam 1:4000) suspended in 10% normal goat serum (NGS). Incubation was performed overnight at 4°C on a shaker. Each rinse was performed for two minutes and repeated three times. Following primary incubation, sections were given three successive two-minute rinses in PBS-Tx. Slices were then incubated in 10% NGS for two hours at room temperature with the secondary antibody, goat anti-rabbit IgG conjugated to the fluorophore Alexa Fluor 647 (Thermo Fisher Scientific 1:200). Slides were then prepared to assess GRIN lens and STN lead placement based on the locations of the hypointense artifact left by removal of the implants before extraction of the brain. GRIN lens placement was visually confirmed to be within the dorsal striatum, a large and easily identifiable target. STN lead placement was confirmed by overlaying of a mouse atlas^52^ on images of each slice showing the hypointense artifact left by the tip of the STN stimulation electrode. Animals were excluded from analysis if neither of the two stimulating electrode contacts were located within the STN as assessed through histological analysis. Following secondary incubation, sections were given three successive two-minute rinses with 0.1M phosphate buffer, mounted onto subbed glass slides, and the coverslip was placed with ProLong Antifade Reagent containing DAPI (Thermo Fischer Scientific). Coronal slices were imaged using the Nikon Eclipse Ti microscope with Lumencor LED light source and Nikon Elements software. For assessment of the 6-OHDA lesion, tyrosine hydroxylase (TH) expression was quantified^61^. Images were obtained with a Tx Red filter cube and a shutter speed of 4ms at 50% LED power. Thresholds were set to (30-255) after performing rolling ball background subtraction with a 50 pixel radius using ImageJ (National Institutes of Health, Bethesda, MD). TH positive cell bodies in the substantia nigra were manually counted as described in Winkler et al.^62^ and Santiago et al.^63^. In the dorsal striatum, mean-pixel intensity of the TH signal was quantified using ImageJ^64^ as the sum of pixel values (0-255)/number of square pixels. Striatal regions deformed by the GRIN lens tract as observed in the histology were not included in the analysis of mean-pixel intensity.

## Results

The goal of this proof-of-principle study was to establish a methodological framework for overcoming experimental and data analysis challenges associated with combining electrical stimulation with calcium imaging in freely moving animals. The experimental protocol consisted of AAV9 injections to induce GCaMP6m expression (Figure 2A), combined with placement of a GRIN lens (Figure 2B), 6-OHDA lesioning (Figure 2A), and placement of bipolar stimulating electrodes (Figure 2B), first validated separately in different animal cohorts.

### Histological Validation of 6-OHDA Lesioning, GCaMP Expression, and Implant Positioning

The full experimental paradigm, consisting of 6-OHDA lesioning, AAV9 injection, GRIN lens implantation and STN stimulating electrode implantation, was performed on 10 mice. Two mice were lost due to surgical complications and two more due to GRIN lens damage that prevented imaging. An additional two mice were excluded from all analyses due to mistargeting of the stimulating electrode outside of the STN as assessed by post-mortem histology. Expression of GCaMP6m calcium indicator, induced by AAV9.CAG.GCaMP6m.WPRE.bGHpA injection, was confirmed via histological analysis that showed green fluorescent cell bodies in the dorsal striatum (Figure 3B). Most GCaMP expressing cells did not show any nuclear expression, which is an important indicator of cell health (Figure S3)^48^. Dopaminergic lesioning via a unilateral 6-OHDA lesion, performed in the ipsilateral hemisphere, was confirmed by immunohistochemical labeling of TH followed by quantification of TH positive cells and mean pixel intensity (Figure 3A). A significant decrease in dorsal striatal mean pixel intensity (Figure 3D, p<0.05; Welch’s unequal variance t-test), and SNc TH positive cells (Figure 3E, p<0.05; Welch’s unequal variance t-test) between the lesioned and non-lesioned hemispheres was observed.

**Figure 3.**
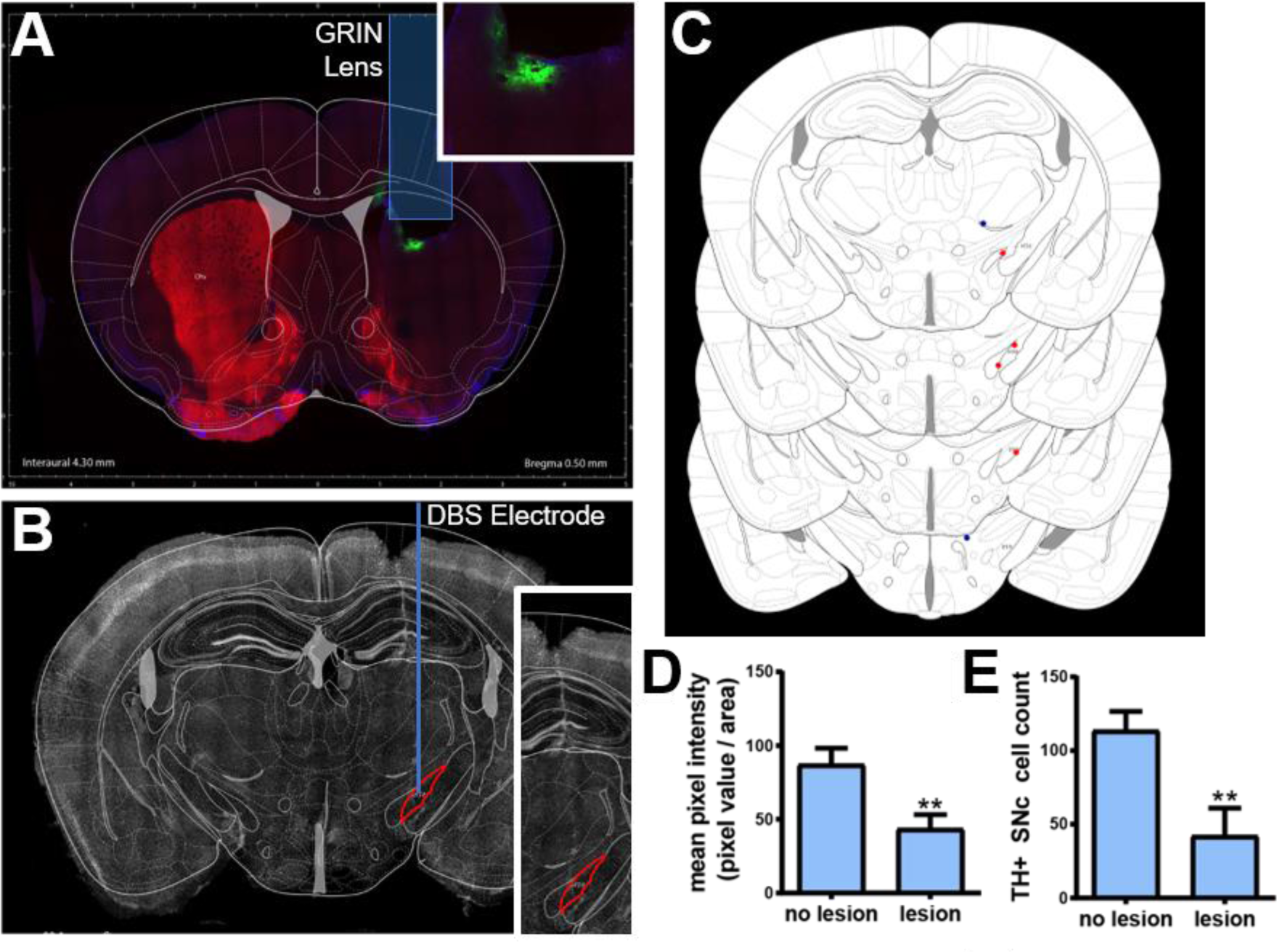
Histological and behavioral assessments. **A)** Coronal view of the histological analysis through the striatum showing lens placement, GCaMP6m expression (green label, inset), tyrosine hydroxylase (TH) stain (red label) demonstrating unilateral dopaminergic lesion in the striatum a healthy animal **B)** Coronal view indicating DBS electrode placement in the subthalamic nucleus (STN). **C)** A coronal atlas view showing DBS lead placement in 6 animals (orange dots indicate electrodes located within the STN, blue dots indicate electrodes placed outside of the STN) compared to the location of the STN. **D)** TH+ mean pixel intensity (0-255) calculated over dorsal striatum, (Welch’s unequal variance t-test, p<.05, n=10) and **E)** quantification of TH positive SNc cells in lesioned and non-lesioned hemispheres (Welch’s unequal variance t-test p<.05, n=12; 1866 total cells), indicating successful 6-OHDA lesioning.

### Chronic Calcium Imaging

Initial validation of calcium imaging within the striatum was performed in healthy mice without stimulation electrode implants (Figure 4). A consistent number of cells, as identified by CNMF-E, were obtained in six recording sessions performed over 33 days across five animals (Figure 4B). This indicates that recordings from the same brain area can be performed over time, however slight differences in the field of view were observed between recording sessions (Figure 4C). Additionally, damage to the microscope during a recording session and the subsequent repair led to a slight change in the focal plane, including a rotation of 90 degrees, between sessions (Figure 4C).

**Figure 4.**
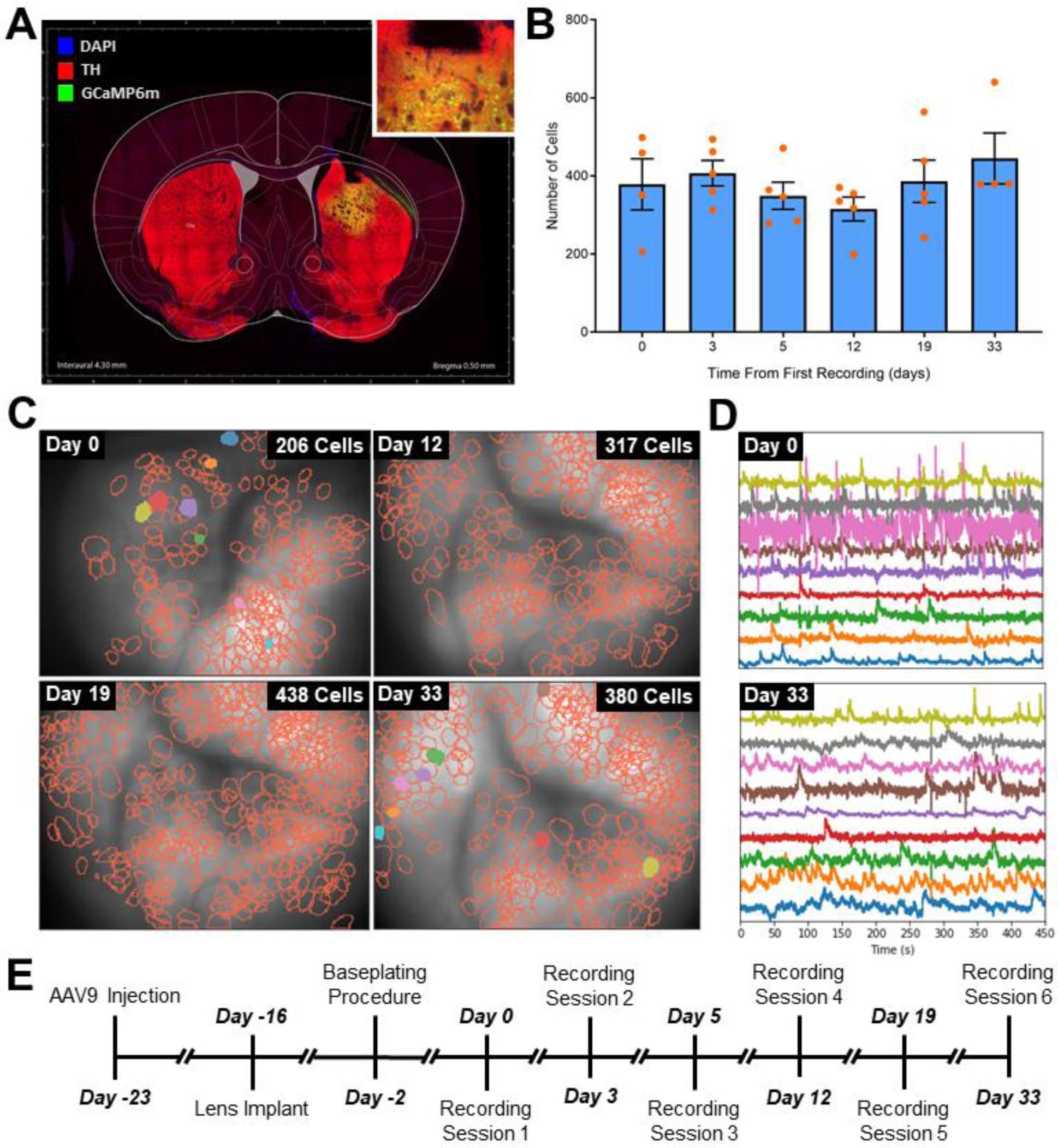
Chronic calcium recordings in healthy mice. **A)** A coronal view through the striatum of a healthy mouse indicating lens placement, GCaMP6m expression (green, inset), but no dopaminergic lesion as shown by the TH+ stain (red) for dopaminergic cells. **B)** The number of cells recorded across a cohort of five mice and identified using constrained non-negative matrix factorization for microendoscope data (CNMF-E) was consistent over 33 days (p=0.89; one-way ANOVA). **C)** Representative mouse showing that the number of striatal neurons identified by CNMF-E over the course of the study (note the rotation of the microscope field of view between days 0 and 12). **D)** Randomly selected calcium traces from 10 cells extracted by CNMF-E. **E)** Experimental timeline for these experiments.

### Neural Signal Identification

Initial validation of the ability to record STN stimulation evoked responses in striatum prior to awake behaving experiments was performed in six anesthetized animals. Two animals did not exhibit stimulation evoked response within striatum; it was later verified via post-mortem histology that neither of the electrodes of the twisted bipolar pair were located in STN, and these animals were removed from the analysis. Stimulation evoked responses were observed in four anesthetized animals that were imaged during 10 second epochs of 30, 80 and 130 Hz stimulation. These epochs were separated by one-minute intervals to eliminate the residual effects from previous stimulations. This sequence was repeated three times in each animal. The striatum was quiescent while stimulation was turned off (Figure 5A) under anesthesia, and only became active when STN stimulation was applied (Figure 5B-D). STN stimulation activated a subset of in-focus cells within the striatum as well as caused out-of-focus changes in background fluorescence (Figure 5B-D), which may originate from stimulation-evoked broad changes in cell activation. These stimulation-dependent out-of-focus signals are exemplified by changes in mean fluorescence within the field-of-view shown in Figure 5E. These out-of-focus changes and synchronous activation of multiple neurons confounded the analysis of calcium signals via standard data analysis techniques such as PCA/ICA, which identifies cells using differences in temporal characteristics (Figure S5B). To quickly assess the performance of common algorithms in isolating activity of individual cells when in the presence of significant out of plane calcium activity induced by stimulation, simulated changes in background fluorescence based on a model of out-of-plane activation were added to an experimental data set (Fig. S2). The capability of common data analysis methods and spatial filtering techniques to identify spatially compact neurons and remove the simulated contamination were then qualitatively compared (Figure S5).

**Figure 5.**
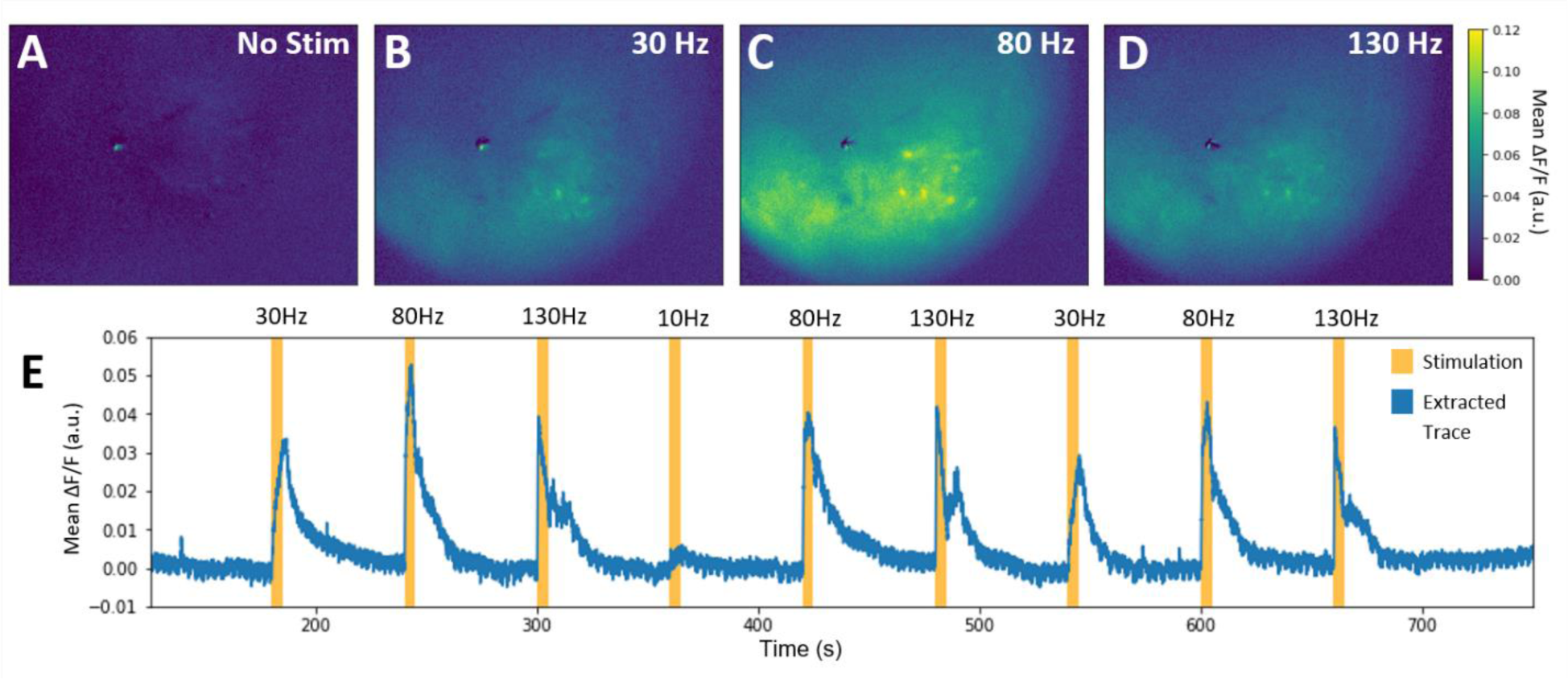
Subthalamic nucleus (STN) stimulation in anesthetized mice.. **A-D)** Representative calcium responses evoked by different stimualation frequencies (no stimulation (A), 30 Hz (B), 80 Hz (C), and 130Hz (D) stimulation) in an anesthetized 6-OHDA lesioned mouse. **E)** The mean flourescence across the field of view during a sequence of stimulations.

ROI analysis, PCA/ICA with or without spatiotemporal de-mixing, and CNMF-E^55,56^ were evaluated for their ability to extract well-defined cell bodies in the presence of real and simulated background fluorescence. Although ROI analysis allows for selection of well-defined cell bodies, it is a subjective technique and cannot separate cell activity from low spatial frequency background contamination. As a result, calcium traces extracted during electrical stimulation of the STN were nearly identical for all cells in the field of view (Figure S5A). This suggests that the traces were primarily the results of background contamination and not from individual neural activity. Additionally, ROI analysis was not able to remove simulated background contamination (Figure S5A). These results suggest that ROI analysis is not well suited for neuron identification from recordings obtained during electrical stimulation. Similar to ROI analysis, PCA/ICA^67^ failed at uniquely identifying cells that exhibited synchronous firing during stimulation and was unable to completely remove artificial background contamination (Figure S5B). Spatiotemporal de-mixing, which can be used to improve the identification of cells with PCA/ICA^67^, did not result in significant improvements (Figure S5C). Both ROI analysis and PCA/ICA were applied to the same data prefiltered using a spatial bandpass filter with cutoff frequencies chosen based on the size of cell bodies observed in the ΔF/F data (5 and 31 pixels). In both cases spatial filtering improved the rejection of artificial background contamination but did not completely eliminate the background contamination (Figure S5D-F). In contrast, CNMF-E was able to identify individual cell bodies despite synchronous neural activity (Figure S5G), does so in an objective manner, and is much less susceptible to low spatial frequency background noise. For this reason, we used CNMF-E for all subsequent data analysis presented in this manuscript.

### Frequency Dependence of STN Electrical Stimulation in Anesthetized Animals

A preliminary characterization of frequency dependent changes in STN stimulation evoked calcium responses within striatum was performed on recordings from four anesthetized mice with stimulation electrodes located in the STN that were described in the previous section. Although the striatum was observed to be quiescent under anesthesia (Figure 5A), clear calcium activity suggestive of neuronal activation occurred with onset of STN stimulation at 30, 80, and 130Hz (Figure 5B-D, Video S1). Additionally, the recruitment occurred in a frequency-dependent manner (Fig. 6A-C) with 80 and 130 Hz stimulation exhibiting a greater number of activated cells than stimulation at 30Hz (Figure 6D, p<0.05; Kruskal-Wallace one-way analysis of variance, Dunn’s multiple comparison test). Despite the overall frequency dependence of neural activation, there was heterogeneity in the response of individual neurons to different stimulations as shown by Figure 6E. For instance, some cells only responded to stimulation at 80 Hz, but not 130Hz (i.e. Figure 6E, cell 2) or even showed heterogeneous responses to repeated stimulation at the same parameters (i.e. Figure 6E, cell 4). The stimulation evoked behavior of specific cells did not necessarily depend on relative location of the cell in the field of view, with co-localized neurons showing different responses to similar parameters (Figure 6E).

**Figure 6.**
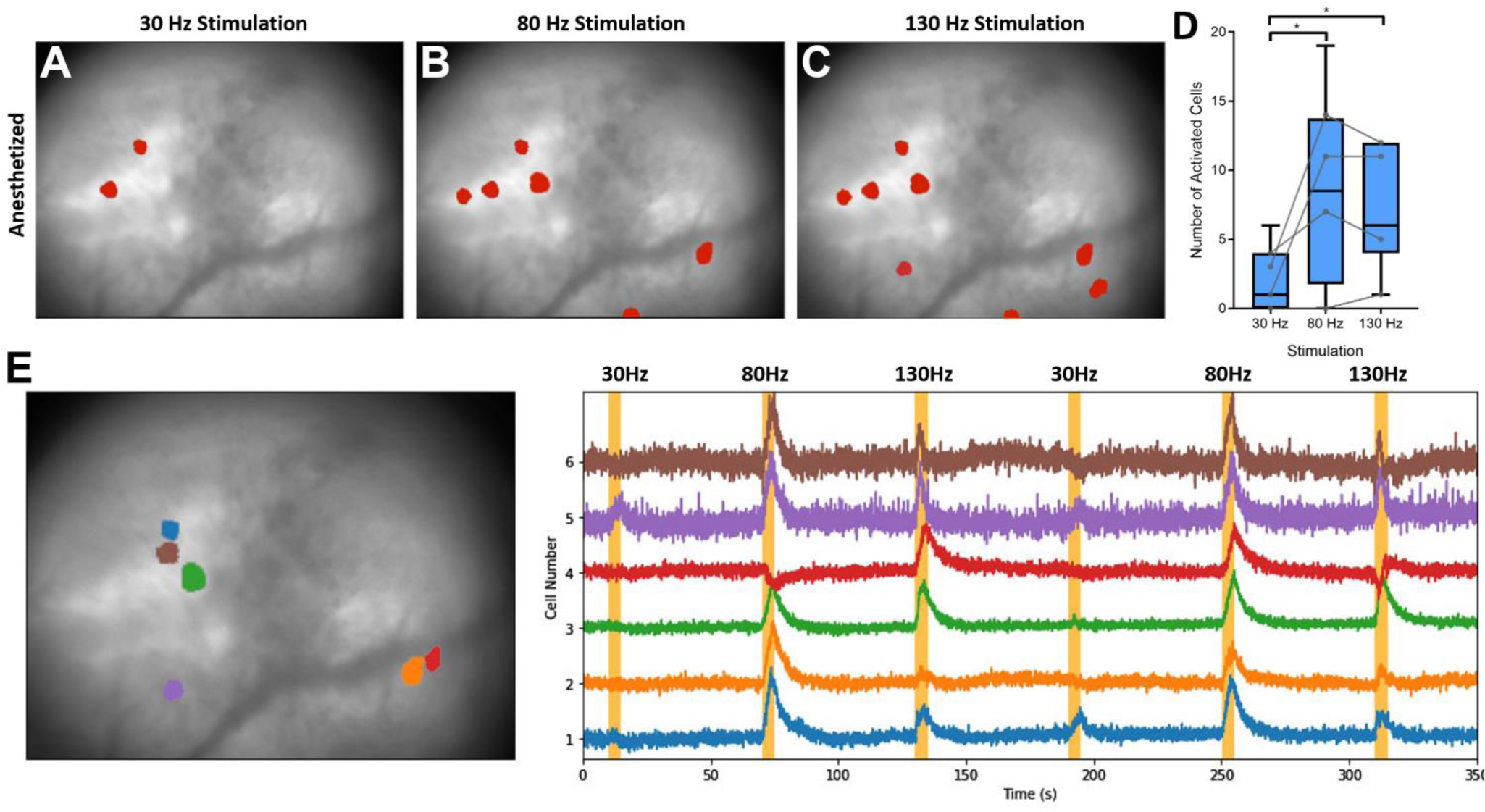
Frequency dependent responses to electrical stimulation of the subthalamic nucleus (STN). **A-C)** Cells responding to 30, 80, and 130 Hz stimulation from 3 consecutive stimuli, respectively, overlaid on the mean background image recorded in a representative anesthetized mouse. **D)** Number of activated cells responding to 30-130Hz stimulation in a cohort of four mice with three repetitions of each stimulation. Kruskal-Wallace one-way ANOVA, Dunn’s multiple comparison test; * indicates p<0.05. Gray dots show the median within-animal responses for each animal. Analysis was performed on 19, 4, 7, and 14 total neurons identified during stimulation under anesthesia. **E)** Heterogeneity of neuronal responses to 30, 80, and 130Hz stimuli in one representative 6-OHDA lesioned mouse.

### Stimulation Evoked Behavioral and Calcium Responses

Two of the four mice in which anesthetized stimulation evoked recordings were performed also underwent awake-behaving imaging sessions in an open-field during stimulation (200µs pulsewidth, 30µA amplitude and 200µs pulsewidth, 80µA amplitude). In contrast to anesthetized recordings in the same animal, the striatal neural population was found to be highly active during open field behavior with increased activity during animal movement (Figure 7A, Video S2). This can be seen in the relationship between animal velocity (Figure 7A top row) and cell activity shown on the raster plots (Figure 7A 2nd row) and calcium traces (Figure 7A 3rd row). The raster plot shown in the 2nd row of Figure 7A shows the calcium event timing for all cells in a representative data set and the traces show) and the raster of calcium traces on the 3^rd^ row of Figure 7A shows the calcium activity for the 40 highest SNR cells in the data set. Significant differences in the relationship between the rate of calcium events and animal velocity were observed at both 30 Hz Figure 7B,C; p<0.0001 Kruskal-Wallace one-way analysis of variance, Dunn’s multiple comparison test) and 130 Hz (Figure 7B,C; p<0.0001 Kruskal-Wallace one-way analysis of variance, Dunn’s multiple comparison test).

**Figure 7.**
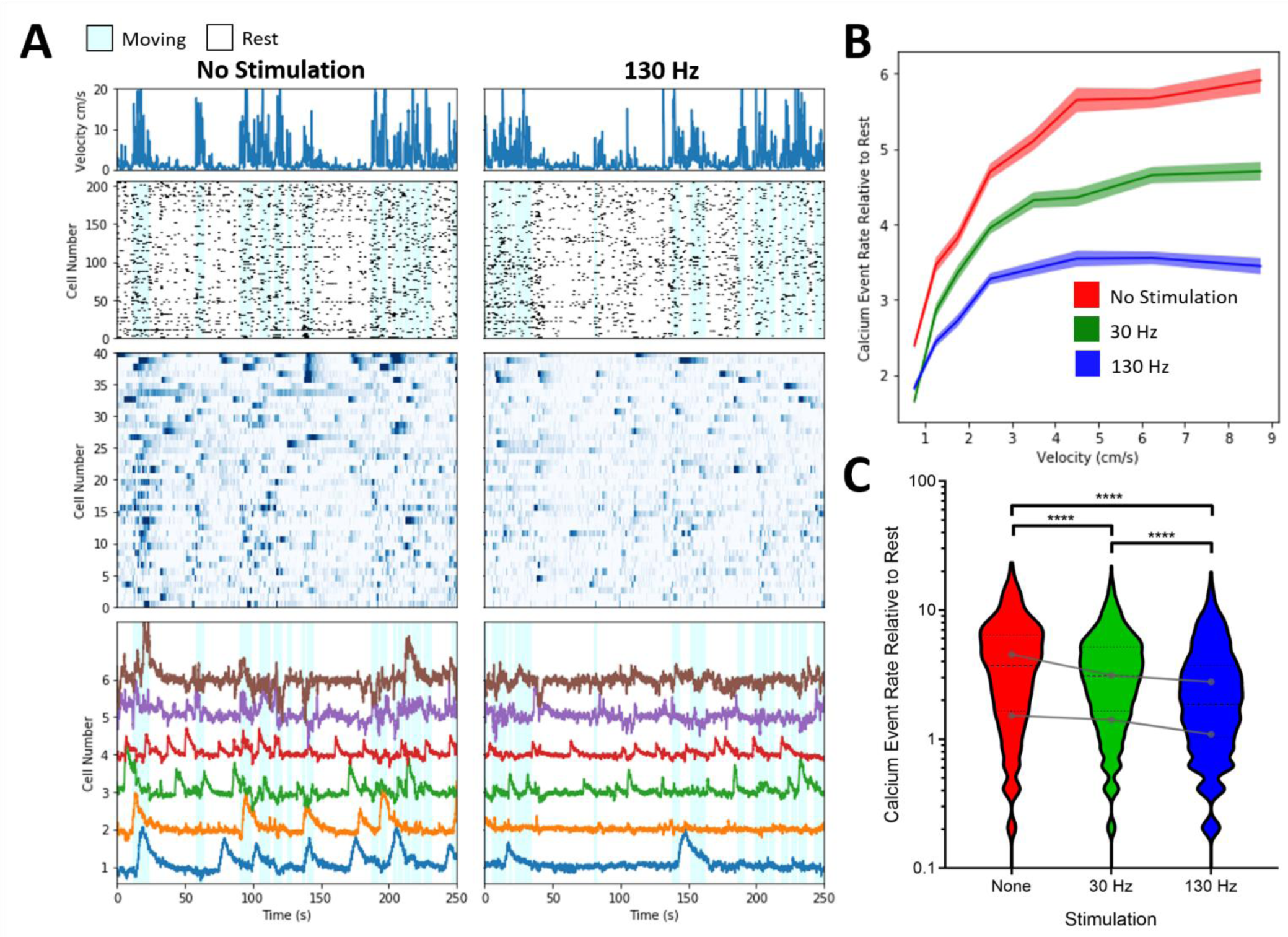
Relationship between neural activity and animal behavior (velocity) change with STN stimulation. **A)** Striatal cell activity (lower panels) identified by constrained non-negative matrix factorization for microendoscope data (CNMF-E) was correlated with mouse behavior (top row), as measured by velocity, across stimulated and unstimulated open field behavior sessions. The second row displays the identified calcium events from all cells in a representative data set, while the third row represents calcium traces from 40 cells with the highest SNR in the same data set. The fourth row shows cell traces for the neurons with identified calcium events that are most correlated with mouse velocity during periods of no stimulation, showing examples increased activity during periods of movement. **B)** Differences in the relationship between the rate of calcium events normalized to no stimulation at rest (velocity < 0.5 cm/s) between all stimulation paradigms. **C)** The rate of calcium events normalized to no stimulation at rest (velocity < 0.5 cm/s) between all stimulation paradigms. Kruskal-Wallace one-way analysis of variance, Dunn’s multiple comparison test; **** indicates p<0.0001. Gray dots show median within animal responses to stimulation. Analysis was performed on 208 and 344 neurons identified in these two animals.

## Discussion

Although there are several advantages to this combination of optical recordings and DBS in behaving pathological animals, achieving the complete experimental paradigm requires successful completion of a complicated set of procedures. All of the experiments outlined to demonstrate the utility of this method require the implementation of four separate components: 1) induction of calcium indicator, GCaMP6m, expression via a unilateral injection of an AAV9 viral vector into striatum (Figure 2A,B), 2) placement of a Gradient Refractive Index (GRIN) lens in the target recording location just above striatum (Figure 2A,C), 3) induction of a Parkinsonian phenotype via unilateral injection of 6-OHDA into the substantia nigra pars compacta (SNc) (Figure 2A,B), and 4) placement of a bipolar stimulating electrode within the STN for application of electrical stimulation (Figure 2A,B). The complexity of troubleshooting these experimental components as well as the need for validation of each component in the context of the complete experimental paradigm is a significant hurdle to the adoption of this method for studying the effects of DBS. With the goal of enabling other researchers to adopt this combination of techniques to the study of neuromodulation therapies, here we describe our approach to implement and validate these experimental components to aid others in performing similar experiments.

### Demonstration of Utility of Calcium Imaging during STN DBS in Parkinsonian Mice

The fundamental goal of the experiments outlined in this methods paper was to demonstrate technical feasibility of the multicomponent surgical procedure, as well as demonstrate that the stimulation evoked responses in a commonly explored downstream structure (striatum) of a traditional DBS target (STN) varied as a function of behavioral state (anesthetized, awake sedentary, awake moving). These data were intended to underscore that the combined experimental preparation can yield unique information that may be useful in the study of DBS mechanisms in the future, but were not intended at this time to make any definitive conclusions about the therapeutic mechanisms of STN DBS to treat parkinsonian symptoms.

As a first exploration of the capabilities of single photon calcium imaging to assess the effects of DBS, we looked at the frequency dependence of evoked neural activity in anesthetized mice. Other studies have shown that electrical stimulation excites neural elements in a frequency dependent manner^68^ and that therapeutic responses to neuromodulation are highly dependent on stimulation frequency^69^. For example, therapeutic efficacy for STN DBS in human patients is typically achieved with high frequency stimulation in the range of 120-180 Hz^70^. The population of neurons activated by stimulation during anesthesia was a small subset of the neurons identified during unstimulated awake behaving recordings (Figure 6A-C). This sparse activation is consistent with other work using optical techniques showing sparse neural activation around a stimulation electrode^68,71^. An increase in the number of cells activated at 80 and 130 Hz stimulation was observed compared to stimulation at 30 Hz (Figure 6D). In anesthetized optical recordings, heterogeneous changes in neural activation within striatum were observed, with different cells that were near each other in the microscope field of view responding to stimulation at different frequencies (Figure 6E). This heterogeneity may be due to differences in response of specific cell types. The heterogeneity could also be the result of subtle changes in stimulation evoked local neuronal/non-neuronal activity as a function of distance from the bipolar electrode, leading to varied responses at a distal point in the connected circuit.

To illustrate the technical feasibility of performing optical recordings in striatum while stimulating STN in an awake behaving pathological model, we performed a series of recordings with 10-minute periods of 30Hz and 130Hz electrical stimulation interleaved by 10-minute periods without stimulation. In behaving mice, the striatum was found to be much more active (Figure 7A) than under anesthesia (Figure 5A). Additionally, neuronal activity within the striatum was found to increase with movement of the mouse, as assessed by animal velocity (Figure 7A-B). In contrast to STN stimulation in the anesthetized state which increased observed activity in striatum (Figure 6), in the two animals in which anesthetized and awake recordings were performed STN stimulation during minimally constrained behavior decreased the activity in striatum (Figure 7). Given the low number of animals we cannot draw any definitive conclusions about DBS mechanism across an entire population; however, these data suggest that the downstream effects of DBS may also depend on behavioral inputs to the same circuitry, thus underscoring the need for recording technology compatible with unrestricted behavioral experiments during DBS. Additionally, recording from genetically defined sub-populations of cells using calcium imaging would enable studies of neuromodulation effects in the context of specific neuronal cell-types, such as D1 and D2 medium spiny neurons that form the primary outputs of the striatum^43^.

### Technical Challenges Toward Implementation of the Complete Experimental Protocol

The complex nature of these proof of concept experiments led to successful stimulation and imaging in 40% of the animals, with primary failure points being anesthesia related complications during surgery (20%), lens damage during group housing of the animals (20%), and failure to place the STN stimulation electrode within the target structure (20%). These failure points can be improved with practice and some simple modifications to the experimental procedures. However, with multiple cumulative points of failure across different components of this procedure, it is particularly important to minimize failures in order to obtain a practical level of experimental yield. To accomplish this, we recommend optimizing each component of the surgical procedure in isolation to minimize surgical time and complications prior to attempting the components in combination. By optimizing each component in parallel, we were able to complete the first cohort of animals with the entire experimental paradigm approximately two months after beginning this project.

The DBS implant procedure has a high rate of failure due to the difficulty targeting small brain nuclei in the mouse mode. For this reason, it is important to practice and optimize the targeting of the DBS implant before combining it with the other components of the overall procedure. Since, validation of electrode location cannot be obtained until histological verification is performed, a large amount of time could be spent performing experiments on animals that will be excluded from the analysis due to mistargeting of the electrode. One step that we have since taken to improve targeting is switching to the use of custom electrodes made from platinum/iridium wire (A-M Systems) instead of the platinum electrodes from PlasticsOne used in these studies. At the electrode sizes used in these experiments (75µm diameter wire), platinum/iridium electrodes are much stiffer than pure platinum electrodes; in our experience they were less likely to deviate from the target trajectory during insertion. Another component that should be independently validated before attempting implantation of the GRIN lens and subsequent imaging is that GCaMP expression in the target location is sufficient and does not show nuclear expression that is indicative of toxicity^48^. Appropriate GCaMP expression is a critical aspect of successful imaging that can be easily checked before imaging in genetic lines or with AAV induced models. If using an AAV, expression can be adjusted by adjusting the titer of the virus, which may be helpful for optimizing for a specific target location.

An avoidable point of failure in our experiments that can be addressed during the surgical procedures was protection of the GRIN lens from damage during the time between implantation of the lens and baseplating the animal. During the experiments presented in this manuscript, we protected the GRIN lens with a protective plastic cap made from the end of a microcentrifuge tube and attached with Kwik-Cast adhesive. We have since switched to using dental cement to hold the protective cap to the animals headcap for a more secure connection. Whatever method is selected, it should be optimized prior to attempting the complete experimental paradigm to prevent animal losses at this stage.

The implementation of microendoscope optical recordings and simultaneous stimulation through implanted electrodes needs to be carefully planned in the context of specific behavioral experiments, as the connection to instrumentation can both constrain behavior as well as place strain on the connectors and the headcap itself. In our experiments, for instance, individual recording sessions were limited to 10 minutes in 6-OHDA lesioned mice in order to prevent tangling of the microscope and stimulation wires due to biased rotational behavior of the lesioned animals. In the future this recording time will be extended with the addition of a commutator^72–74^. Appropriate control of the wires and use of a commutator to enable unrestricted behavior of the animal will be critical to rigorous studies of behavioral changes during the application of DBS, where additional wires needed for electrical stimulation can impose additional constraints on behavior of the mouse over the miniature microscope alone.

Other studies using single and two photon microscopy in behaving animals have shown stable longitudinal recordings in excess of one month^43–46^. These data suggest that longitudinal recording from the same cell population over weeks of neuromodulation therapy may be possible. Neuromodulation therapies for some neurologic and psychiatric conditions, such as depression and epilepsy can take months to show therapeutic effect^11,34–42^. Thus, the capability to longitudinally record from the same set of cells via calcium imaging in a deep brain regions over weeks to months is important to enable the *in vivo* study of plasticity in neuromodulation therapies that can be difficult via electrophysiology or functional imaging methods. In contrast to electrophysiological recordings where highly similar waveforms measured across time from different neurons in the vicinity of the recording electrode can make it difficult to guarantee the tracking of a single-unit from recording obtained on different days^24,25^, the ability to directly visualize neuron somas in optical recordings and use their relative locations as landmarks may make it easier to confidently track the same neuron over time^66^. In Figure 4B, we demonstrated the ability to record cells during 6 recording sessions over the course of a month. However, slight differences in the field of view can be seen between recording sessions as well as a rotation of the field of view caused by the repair of the microscope being used in the experiments between days 0 and 12. We attempted to perform tracking of individual cells between recording session in the data from these experiments using the cell tracking algorithm presented in Sheintuch et al. 2018. However, this was not successful due to inconsistent placement of the microscope leading to changes in the field of view between sessions. For this reason, although it is likely possible to align cells over longer periods of time up to weeks or months^43–46^ in animals undergoing neuromodulation therapies, it is important to note that these analysis require careful placement of the microscope to obtain the same field of view between recording sessions. In practice this proved to be difficult, however the process may be improved by new iterations of microscope technology that include electronic focusing capabilities^75^.

### Neural Signal Identification Techniques during Electrical Stimulation

Unlike two-photon microscopy, which is robust to changes in activity outside of the focal plane, single photon microscopy can be easily contaminated by out-of-focus changes in neural activity. Although these changes in activity may not significantly impact the analysis of neural activity during some behavioral paradigms, they can have a significant effect when electrical stimulation is applied. Therefore, attention should be paid to application of existing analysis techniques to the study of neuromodulation paradigms, which may introduce non-physiological changes in neural activity. Initially, we applied PCA/ICA, a commonly used analysis technique for calcium imaging data, to identify cell activity in recordings obtained in behaving animals. Using PCA/ICA we observed artifactual high correlation between calcium traces recorded from nearby cells and were not able to distinguish overlapping cells bodies (Figure S4), a limitation that have been observed by other recent studies.^55,65^ This led us to perform a qualitative analysis of different algorithms to remove artificially added stimulation induced background fluorescence changes from data recorded in anesthetized animals (Figure S5). Our data suggests that ROI analysis and PCA/ICA, which are commonly applied algorithms for identifying neural signals in calcium imaging recordings, were not suitable for isolating calcium traces from in focus neurons from the artificially added out-of-focus contamination. Compared to ROI analysis and PCA/ICA, even with bandpass filtering of the data prior to analysis, CNMF-E was better able to isolate calcium traces from in focus neurons. This suggests that CNMF-E is better suited analysis of data obtained during the application of neuromodulation therapies. These results is in agreement with other comparisons of CNMF-E to PCA/ICA in other contexts that have been recently published^55,65^.

### Validation of the Pathological Model and DBS Therapy in the Context of the GRIN Lens Implant

In order to perform a mechanistic study to isolate the mechanisms of STN DBS therapy with appropriate scientific rigor in the mouse model, there are a number of potential confounds that must be addressed, including: 1) establishing the therapeutic efficacy of DBS to alleviate pathological symptoms in the mouse model without the complication of an invasive indwelling GRIN lens, 2) determining the impact of the indwelling GRIN lens on normal and pathological behavior, and 3) assesing the impact of the indwelling GRIN lens on the therapeutic function of DBS.

Single-photon calcium imaging via a head-mounted microscope is primarily performed in mouse models due to the availability of genetic lines and more extensive experience with induction of expression with viral vectors that can be leveraged to express calcium indicators in specific cell types. In contrast, historic studies of neuromodulation therapies usually utilize rats or large animal models in order to increase space for the implantation of electrodes, improve targeting of small neural structures, and reduce the difference in sizes of electrodes and brain nuclei compared to the human therapies. Although many pathological models for diseases treated by DBS have been well established in mouse models, the electrode configuration and parameters that best mimic clinical DBS in mouse animal models are not well understood.^49,76^ Therefore, the therapeutic efficacy of DBS applied to clinically relevant targets in these models needs to be validated before rigorous studies into the mechanisms of DBS can be performed. We have begun this work for STN-DBS in a 6-OHDA mouse model of Parkinson’s Disease, with preliminary data provided in the Supplemental Methods as a frame of reference (Figure S2).

Another important consideration that needs to be further validated is the effect of tissue aspiration and GRIN lens implantation on the mechanisms underlying the effects of neuromodulation therapies^77^. For example, this procedure disrupts tissue within the motor cortex, striatum, and other structures which may be critical to the therapeutic effects of DBS. There is growing evidence that the trauma due to the implantation of electrodes in or on the brain, and the trauma due to the associated surgical procedure, can induce deficits in motor function and other outcomes. These deficits may only be apparent when more sophisticated measures of motor function are used^77^. Similarly, there is evidence to suggest the presence of a comparatively small chronic indwelling electrode may impact ion channel expression in nearby tissue out as far as 200 microns^78^. For this reason, it is paramount to understand and anticipate potential confounds created by the GRIN lens implant on both normal and pathological behaviors, as well as response to therapeutic interventions. To some extent the lens implant access route can be optimized to minimize influence on pathways putatively associated with the behavior in question, pathological influence on that behavior, and circuitry hypothesized to be engaged in DBS therapeutic effect. DBS therapies are increasingly thought to act through circuit mechanisms that rely on activation of tracts of passage near the electrode in addition to local effects. For instance, one of the current leading theories for the mechanism of STN DBS to treat PD is that DBS disrupts pathological synchrony in the basal ganglia-thalamo-cortical motor circuit through antidromic activation of motor cortex^79,80^. In general, disruption of these pathways due to damage caused by aspiration and insertion of the GRIN lens has the potential to disrupt or modify the pathological behaviors being studied or the therapeutic mechanism of DBS.

To address these issues for the 6-OHDA mouse model of Parkinson’s Disease cohorts directly comparing normal behavior with and without a GRIN lens implant in the target recording area, as well as comparing DBS effects on pathological behavior in animal cohorts with and without a GRIN lens implant, must be completed in future studies. It is important to note that changing the route of access for the GRIN lens intended to image the same target could alter pathological behavior and DBS therapeutic efficacy; therefore, any change to the GRIN lens implantation route would need to be assessed with additional cohorts.

## Conclusions

Optical imaging technologies such as single and two-photon imaging of calcium indicators as surrogates of neural activity have been widely adopted by neuroscientists for the ability of these systems to record from hundreds of single cells with cell-type specificity provided by genetically encoded indicators. These techniques are particularly well suited for studying the spatial dynamics in neural activity that orchestrate healthy behavior and the transition to maladaptive dynamics in a disease state^43^. Single photon microscopy via head-mounted microscopes has the distinct advantage of enabling calcium imaging in freely behaving animals and has already been utilized to show how cell-type specific spatial and temporal dynamics within the striatum break down in PD and are restored with pharmacological treatment^43^. Although this study describes preliminary data on the application of optical imaging, and specifically head-mounted miniature microscopy, for the study of electrical stimulation of the STN, these data represent only one example of the myriad of applications for this technology in the study of the neuronal effects of neuromodulation.

## Author Contributions

JKT, AJA, ENN, JLL, and KAL, designed the experiments. JKT, AJA, ENN, and DC performed the experiments. JKT, AJA, ENN, JMT, and NAK analyzed the data. JKT, AJA, ENN, JLL, and KAL wrote the manuscript. TDK, MS, JJN, SLO, JGP, provided additional materials or expertise needed to perform the experiments or analyze the data. JLL and KAL supervised all aspects of the work.

## Acknowledgements

This work was supported by the National Institutes of Health NINDS R01 NS084975, and The Grainger Foundation.

## References

1. Organization, W. H. Neurological disorders. Public health challenges. (World Health Organization, 2007).

2. Al-Harbi, K. S. Treatment-resistant depression: therapeutic trends, challenges, and future directions. Patient Prefer. Adherence 6, 369–388 (2012).

3. Laxer, K. D. et al. The consequences of refractory epilepsy and its treatment. Epilepsy Behav. 37, 59–70 (2014).

4. Benabid, A. L., Pollak, P., Louveau, A., Henry, S. & de Rougemont, J. Combined (thalamotomy and stimulation) stereotactic surgery of the VIM thalamic nucleus for bilateral Parkinson disease. Appl. Neurophysiol. 50, 344–346 (1987).

5. Benabid, A. L. et al. Long-term suppression of tremor by chronic stimulation of the ventral intermediate thalamic nucleus. The Lancet 337, 403–406 (1991).

6. Blomstedt, P. & Hariz, M. I. Deep brain stimulation for movement disorders before DBS for movement disorders. Parkinsonism Relat. Disord. 16, 429–433 (2010).

7. Tröster, A. I., Meador, K. J., Irwin, C. P. & Fisher, R. S. Memory and mood outcomes after anterior thalamic stimulation for refractory partial epilepsy. Seizure 45, 133–141 (2017).

8. Salanova, V. et al. Long-term efficacy and safety of thalamic stimulation for drug-resistant partial epilepsy. Neurology 84, 1017–1025 (2015).

9. Fisher, R. et al. Electrical stimulation of the anterior nucleus of thalamus for treatment of refractory epilepsy. Epilepsia 51, 899–908 (2010).

10. Thomas, G. P. & Jobst, B. C. Critical review of the responsive neurostimulator system for epilepsy. Med. Devices Auckl. NZ 8, 405–411 (2015).

11. Mayberg, H. S. et al. Deep brain stimulation for treatment-resistant depression. Neuron 45, 651–660 (2005).

12. Min, H.-K. et al. Subthalamic Nucleus Deep Brain Stimulation Induces Motor Network BOLD Activation: Use of a High Precision MRI Guided Stereotactic System for Nonhuman Primates. Brain Stimulat. 7, 603–607 (2014).

13. Phillips, M. D. et al. Parkinson Disease: Pattern of Functional MR Imaging Activation during Deep Brain Stimulation of Subthalamic Nucleus—Initial Experience. Radiology 239, 209–216 (2006).

14. Shon, Y.-M. et al. Effect of Chronic Deep Brain Stimulation of the Subthalamic Nucleus for Frontal Lobe Epilepsy: Subtraction SPECT Analysis. Stereotact. Funct. Neurosurg. 83, 84–90 (2005).

15. Benazzouz, A. et al. Effect of high-frequency stimulation of the subthalamic nucleus on the neuronal activities of the substantia nigra pars reticulata and ventrolateral nucleus of the thalamus in the rat. Neuroscience 99, 289–295 (2000).

16. Hashimoto, T., Elder, C. M., Okun, M. S., Patrick, S. K. & Vitek, J. L. Stimulation of the Subthalamic Nucleus Changes the Firing Pattern of Pallidal Neurons. J. Neurosci. 23, 1916–1923 (2003).

17. Kita, H., Tachibana, Y., Nambu, A. & Chiken, S. Balance of Monosynaptic Excitatory and Disynaptic Inhibitory Responses of the Globus Pallidus Induced after Stimulation of the Subthalamic Nucleus in the Monkey. J. Neurosci. 25, 8611–8619 (2005).

18. Maurice, N., Thierry, A.-M., Glowinski, J. & Deniau, J.-M. Spontaneous and Evoked Activity of Substantia Nigra Pars Reticulata Neurons during High-Frequency Stimulation of the Subthalamic Nucleus. J. Neurosci. 23, 9929–9936 (2003).

19. Miocinovic, S. et al. Computational Analysis of Subthalamic Nucleus and Lenticular Fasciculus Activation During Therapeutic Deep Brain Stimulation. J. Neurophysiol. 96, 1569–1580 (2006).

20. Smith, I. D. & Grace, A. A. Role of the subthalamic nucleus in the regulation of nigral dopamine neuron activity. Synap. N. Y. N 12, 287–303 (1992).

21. Jonckers, E., Shah, D., Hamaide, J., Verhoye, M. & Van der Linden, A. The power of using functional fMRI on small rodents to study brain pharmacology and disease. Front. Pharmacol. 6, (2015).

22. Lancelot, S. & Zimmer, L. Small-animal positron emission tomography as a tool for neuropharmacology. Trends Pharmacol. Sci. 31, 411–417 (2010).

23. Lewis, C., Bosman, C. & Fries, P. Recording of brain activity across spatial scales. Curr. Opin. Neurobiol. 32, 68–77 (2015).

24. Dickey, A. S., Suminski, A., Amit, Y. & Hatsopoulos, N. G. Single-Unit Stability Using Chronically Implanted Multielectrode Arrays. J. Neurophysiol. 102, 1331–1339 (2009).

25. Tolias, A. S. et al. Recording chronically from the same neurons in awake, behaving primates. J. Neurophysiol. 98, 3780–3790 (2007).

26. Parastarfeizabadi, M. & Kouzani, A. Z. Advances in closed-loop deep brain stimulation devices. J. NeuroEngineering Rehabil. 14, 79 (2017).

27. Nicolai, E. et al. Design Choices for Next-Generation Neurotechnology Can Impact Motion Artifact in Electrophysiological and Fast-Scan Cyclic Voltammetry Measurements. Micromachines 9, 494 (2018).

28. Polikov, V. S., Tresco, P. A. & Reichert, W. M. Response of brain tissue to chronically implanted neural electrodes. J. Neurosci. Methods 148, 1–18 (2005).

29. Butson, C. R., Maks, C. B. & McIntyre, C. C. Sources and effects of electrode impedance during deep brain stimulation. Clin. Neurophysiol. Off. J. Int. Fed. Clin. Neurophysiol. 117, 447–454 (2006).

30. Fenoy, A. J., Goetz, L., Chabardès, S. & Xia, Y. Deep brain stimulation: are astrocytes a key driver behind the scene? CNS Neurosci. Ther. 20, 191–201 (2014).

31. Ni, Z. et al. Pallidal deep brain stimulation modulates cortical excitability and plasticity. Ann. Neurol. 83, 352–362 (2018).

32. So, R. Q., McConnell, G. C. & Grill, W. M. Frequency-dependent, transient effects of subthalamic nucleus deep brain stimulation on methamphetamine-induced circling and neuronal activity in the hemiparkinsonian rat. Behav. Brain Res. 320, 119–127 (2017).

33. Giannicola, G. et al. The effects of levodopa and ongoing deep brain stimulation on subthalamic beta oscillations in Parkinson’s disease. Exp. Neurol. 226, 120–127 (2010).

34. Broadway, J. M. et al. Frontal Theta Cordance Predicts 6-Month Antidepressant Response to Subcallosal Cingulate Deep Brain Stimulation for Treatment-Resistant Depression: A Pilot Study. Neuropsychopharmacology 37, 1764–1772 (2012).

35. Schulze-Bonhage, A. Brain stimulation as a neuromodulatory epilepsy therapy. Seizure 44, 169–175 (2017).

36. Agnesi, F., Johnson, M. D. & Vitek, J. L. Deep brain stimulation: how does it work? Handb. Clin. Neurol. 116, 39–54 (2013).

37. Temperli, P. et al. How do parkinsonian signs return after discontinuation of subthalamic DBS? Neurology 60, 78–81 (2003).

38. Servello, D., Porta, M., Sassi, M., Brambilla, A. & Robertson, M. M. Deep brain stimulation in 18 patients with severe Gilles de la Tourette syndrome refractory to treatment: the surgery and stimulation. J. Neurol. Neurosurg. Psychiatry 79, 136–142 (2008).

39. Sachdev, P. S. et al. Deep brain stimulation of the antero-medial globus pallidus interna for Tourette syndrome. PloS One 9, e104926 (2014).

40. Greenberg, B. D. et al. Deep brain stimulation of the ventral internal capsule/ventral striatum for obsessive-compulsive disorder: worldwide experience. Mol. Psychiatry 15, 64–79 (2010).

41. Tierney, T. S., Abd-El-Barr, M. M., Stanford, A. D., Foote, K. D. & Okun, M. S. Deep brain stimulation and ablation for obsessive compulsive disorder: evolution of contemporary indications, targets and techniques. Int. J. Neurosci. 124, 394–402 (2014).

42. Dalkilic, E. B. Neurostimulation Devices Used in Treatment of Epilepsy. Curr. Treat. Options Neurol. 19, 7 (2017).

43. Parker, J. G. et al. Diametric neural ensemble dynamics in parkinsonian and dyskinetic states. Nature 557, 177–182 (2018).

44. Barretto, R. P. J. et al. Time-lapse imaging of disease progression in deep brain areas using fluorescence microendoscopy. Nat. Med. 17, 223–228 (2011).

45. Ziv, Y. & Ghosh, K. K. Miniature microscopes for large-scale imaging of neuronal activity in freely behaving rodents. Curr. Opin. Neurobiol. 32, 141–147 (2015).

46. Andermann, M. L. et al. Chronic Cellular Imaging of Entire Cortical Columns in Awake Mice Using Microprisms. Neuron 80, 900–913 (2013).

47. Breese, G. R. & Traylor, T. D. Depletion of brain noradrenaline and dopamine by 6-hydroxydopamine. Br. J. Pharmacol. 42, 88–99 (1971).

48. Resendez, S. L. et al. Visualization of cortical, subcortical and deep brain neural circuit dynamics during naturalistic mammalian behavior with head-mounted microscopes and chronically implanted lenses. Nat. Protoc. 11, 566–597 (2016).

49. Gubellini, P., Salin, P., Kerkerian-Le Goff, L. & Baunez, C. Deep brain stimulation in neurological diseases and experimental models: From molecule to complex behavior. Prog. Neurobiol. 89, 79–123 (2009).

50. Oueslati, A. et al. High-Frequency Stimulation of the Subthalamic Nucleus Potentiates l-DOPA-Induced Neurochemical Changes in the Striatum in a Rat Model of Parkinson’s Disease. J. Neurosci. 27, 2377–2386 (2007).

51. Boulet, S. et al. Subthalamic Stimulation-Induced Forelimb Dyskinesias Are Linked to an Increase in Glutamate Levels in the Substantia Nigra Pars Reticulata. J. Neurosci. 26, 10768–10776 (2006).

52. Paxinos, G. & Franklin, K. B. J. The Mouse Brain in Stereotaxic Coordinates. (Gulf Professional Publishing, 2004).

53. Barbera, G. et al. Spatially Compact Neural Clusters in the Dorsal Striatum Encode Locomotion Relevant Information. Neuron 92, 202–213 (2016).

54. Klaus, A. et al. The Spatiotemporal Organization of the Striatum Encodes Action Space. Neuron 95, 1171–1180.e7 (2017).

55. Zhou, P. et al. Efficient and accurate extraction of in vivo calcium signals from microendoscopic video data. eLife (2018). doi:10.7554/eLife.28728

56. Pnevmatikakis, E. A. et al. Simultaneous Denoising, Deconvolution, and Demixing of Calcium Imaging Data. Neuron 89, 285–299 (2016).

57. Kondo, T. et al. Calcium Transient Dynamics of Neural Ensembles in the Primary Motor Cortex of Naturally Behaving Monkeys. Cell Rep. 24, 2191–2195.e4 (2018).

58. Thévenaz, P., Ruttimann, U. E. & Unser, M. A pyramid approach to subpixel registration based on intensity. IEEE Trans. Image Process. Publ. IEEE Signal Process. Soc. 7, 27–41 (1998).

59. Sivagnanam, S. et al. Introducing The Neuroscience Gateway. CEUR Workshop Proc. 993, 7 (2013).

60. Lempka, S. F., Miocinovic, S., Johnson, M. D., Vitek, J. L. & McIntyre, C. C. In vivo impedance spectroscopy of deep brain stimulation electrodes. J. Neural Eng. 6, 046001 (2009).

61. Iancu, R., Mohapel, P., Brundin, P. & Paul, G. Behavioral characterization of a unilateral 6-OHDA-lesion model of Parkinson’s disease in mice. Behav. Brain Res. 162, 1–10 (2005).

62. Winkler, C., Kirik, D., Björklund, A. & Cenci, M. A. l-DOPA-Induced Dyskinesia in the Intrastriatal 6-Hydroxydopamine Model of Parkinson’s Disease: Relation to Motor and Cellular Parameters of Nigrostriatal Function. Neurobiol. Dis. 10, 165–186 (2002).

63. Santiago, R. M. et al. Depressive-like behaviors alterations induced by intranigral MPTP, 6-OHDA, LPS and rotenone models of Parkinson’s disease are predominantly associated with serotonin and dopamine. Prog. Neuropsychopharmacol. Biol. Psychiatry 34, 1104–1114 (2010).

64. Xavier, L. L. et al. A simple and fast densitometric method for the analysis of tyrosine hydroxylase immunoreactivity in the substantia nigra pars compacta and in the ventral tegmental area. Brain Res. Brain Res. Protoc. 16, 58–64 (2005).

65. Lu, J. et al. MIN1PIPE: A Miniscope 1-Photon-Based Calcium Imaging Signal Extraction Pipeline. Cell Rep. 23, 3673–3684 (2018).

66. Sheintuch, L. et al. Tracking the Same Neurons across Multiple Days in Ca2+ Imaging Data. Cell Rep. 21, 1102–1115 (2017).

67. Mukamel, E. A., Nimmerjahn, A. & Schnitzer, M. J. Automated analysis of cellular signals from large-scale calcium imaging data. Neuron 63, 747–760 (2009).

68. Michelson, N. J., Islam, R., Vazquez, A. L., Ludwig, K. A. & Kozai, T. D. Y. Calcium activation of frequency dependent temporally phasic, localized, and dense population of cortical neurons by continuous electrical stimulation. (2018). doi:10.1101/338525

69. Dostrovsky, J. O. & Lozano, A. M. Mechanisms of deep brain stimulation. Mov. Disord. Off. J. Mov. Disord. Soc. 17 Suppl 3, S63–68 (2002).

70. Grill, W. M., Snyder, A. N. & Miocinovic, S. Deep brain stimulation creates an informational lesion of the stimulated nucleus. Neuroreport 15, 1137–1140 (2004).

71. Histed, M. H., Bonin, V. & Reid, R. C. Direct Activation of Sparse, Distributed Populations of Cortical Neurons by Electrical Microstimulation. Neuron 63, 508–522 (2009).

72. Spieles-Engemann, A. L. et al. Stimulation of the rat subthalamic nucleus is neuroprotective following significant nigral dopamine neuron loss. Neurobiol. Dis. 39, 105–115 (2010).

73. Li, X.-H. et al. High-frequency stimulation of the subthalamic nucleus restores neural and behavioral functions during reaction time task in a rat model of Parkinson’s disease. J. Neurosci. Res. 88, 1510–1521 (2010).

74. Liberti, W. A., Perkins, L. N., Leman, D. P. & Gardner, T. J. An open source, wireless capable miniature microscope system. J. Neural Eng. 14, 045001 (2017).

75. Sato, M. et al. Fast varifocal two-photon microendoscope for imaging neuronal activity in the deep brain. Biomed. Opt. Express 8, 4049–4060 (2017).

76. Halpern, C. H., Attiah, M. A., Tekriwal, A. & Baltuch, G. H. A step-wise approach to deep brain stimulation in mice. Acta Neurochir. (Wien) 156, 1515–1521 (2014).

77. Goss-Varley, M. et al. Microelectrode implantation in motor cortex causes fine motor deficit: Implications on potential considerations to Brain Computer Interfacing and Human Augmentation. Sci. Rep. 7, 15254 (2017).

78. Salatino, J. W., Kale, A. P. & Purcell, E. K. Alterations in ion channel expression surrounding implanted microelectrode arrays in the brain. bioRxiv 518811 (2019). doi:10.1101/518811

79. McIntyre, C. C. & Hahn, P. J. Network Perspectives on the Mechanisms of Deep Brain Stimulation. Neurobiol. Dis. 38, 329–337 (2010).

80. Hammond, C., Ammari, R., Bioulac, B. & Garcia, L. Latest view on the mechanism of action of deep brain stimulation. Mov. Disord. 23, 2111–2121 (2008).

