## Supplementary Information for "Calcium imaging in freely-moving mice during electrical stimulation of deep brain structures"

### Supplementary Materials

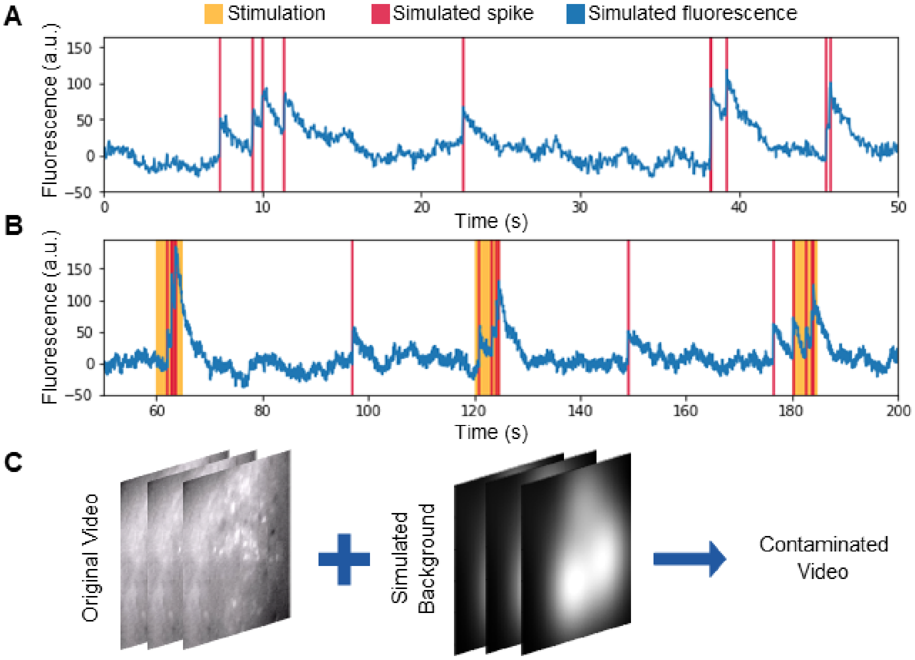


**Figure S1. Method for simulating background florescence contaminated calcium imaging data. A)** Simulation of florescence changes occurring neural activity was performed according to Vogelstein et al. (2009)[68]. **B)** Simulation were performed such that the activity rate of the simulated traces increased during periods of stimulation. **C)** Simulated background fluorescence due to out-of-focus neurons was added to real calcium imaging data obtained under anesthesia to produce a contaminated video that could be analyzed via standard techniques. These data were used to determine the effectiveness of different analysis techniques for identifying cells in the presence of changes in out-of-focus fluorescence occurring during stimulation.

*Behavioral Evaluation of 6-OHDA Induced Parkinsonian Phenotype*

A modified cylinder test was used for the assessment of biased forelimb preference induced by the 6-OHDA lesion[69]. This test is a commonly used method for assessing limb-use asymmetry as a metric for pathologic behavior in 6-OHDA lesioned animals[70–72], Mice were placed in a glass cylinder (15cm wide and 20cm in high) without a prior habituation session and video was recorded 5 min for later scoring by an investigator blinded to treatment group assignments. The number of wall touches (weight bearing paw contact with wall) contralateral and ipsilateral to 6-OHDA lesioned hemisphere were counted during slow-motion playback in VLC media player. These data are presented as a percentage of contralateral touches, calculated as (contralateral touches)/(ipsilateral touches + contralateral touches) × 100.

A cohort of 15 mice underwent 6-OHDA lesioning, implant of a mock GRIN lens (a glass rod of the same dimensions), and implant of a stimulating electrode into the STN. Of these, 8 animals were removed from the study due to failure of the stimulating electrode assessed by electrical impedance spectroscopy and using an impedance value greater than 120 kΩ at 130Hz as a cutoff between failed and working electrodes. A cylinder test, which is a commonly used method for assessing limb-use asymmetry in 6-OHDA lesioned animals[70–72], showed a significant decrease in the proportion of contralateral to ipsilateral paw touches as a result of 6-OHDA lesioning (Figure 3F, $p<0.005$; repeated measures one-way anova with post-hoc tukey test). A non-significant increase in the proportion of contralateral to ipsilateral paw touches was observed in mice electrically stimulated in the ipsilateral STN (Figure 3F, $p=0.053$; repeated measures one-way anova with post-hoc tukey test).

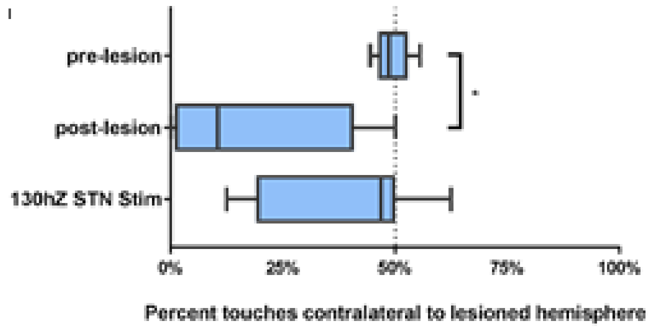


**Figure S2. Behavioral assessment.** Percentage contralateral paw touches pre and post 6-OHDA lesion and with 130Hz STN stimulation ($p<0.005$, Data represented as median, lower/upper quartiles and extremities, repeated measures one-way ANOVA with post-hoc Tukey's test), indicating greater limb-use asymmetry in 6-OHDA lesioned animals.

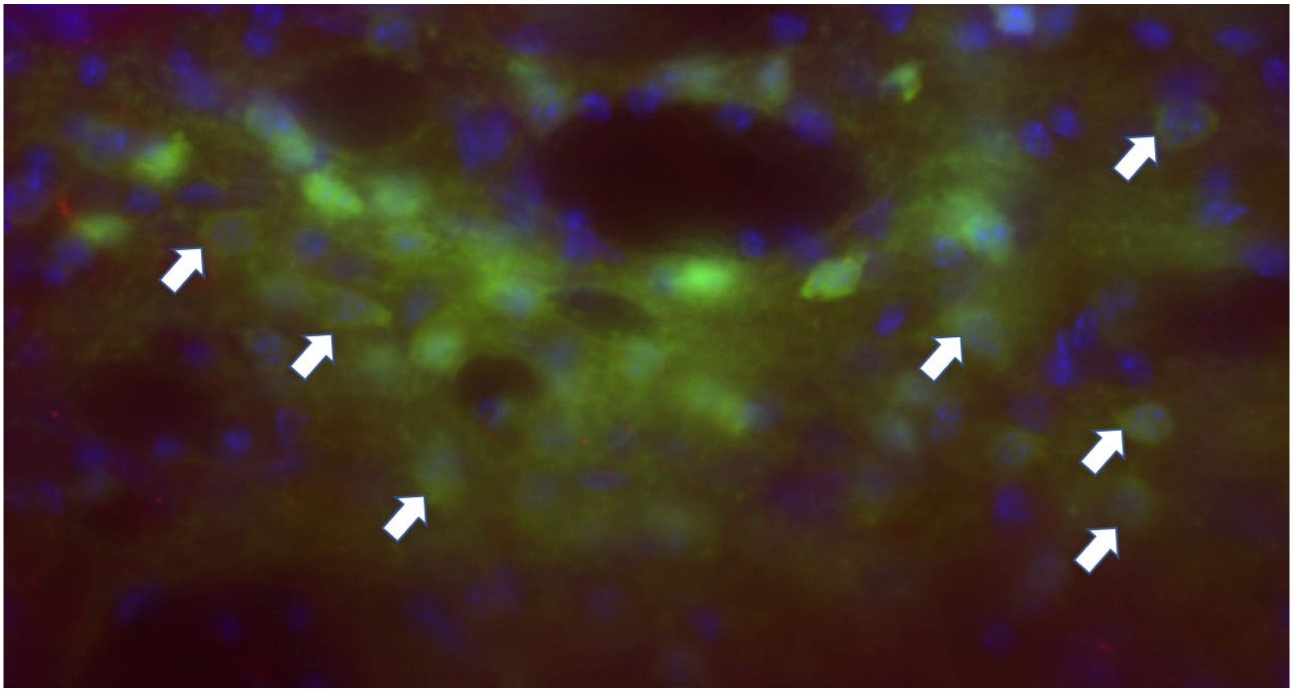

**Figure S3. Histological assessment.** Zoom in of GCaMP6m expression from a section of tissue from the dorsal striatum that just below the lens. This image shows cells with GCaMP expression (green) and without filled nuclei, one commonly used indicator of cell health. Cell nuclei are shown by DAPI stain (blue).

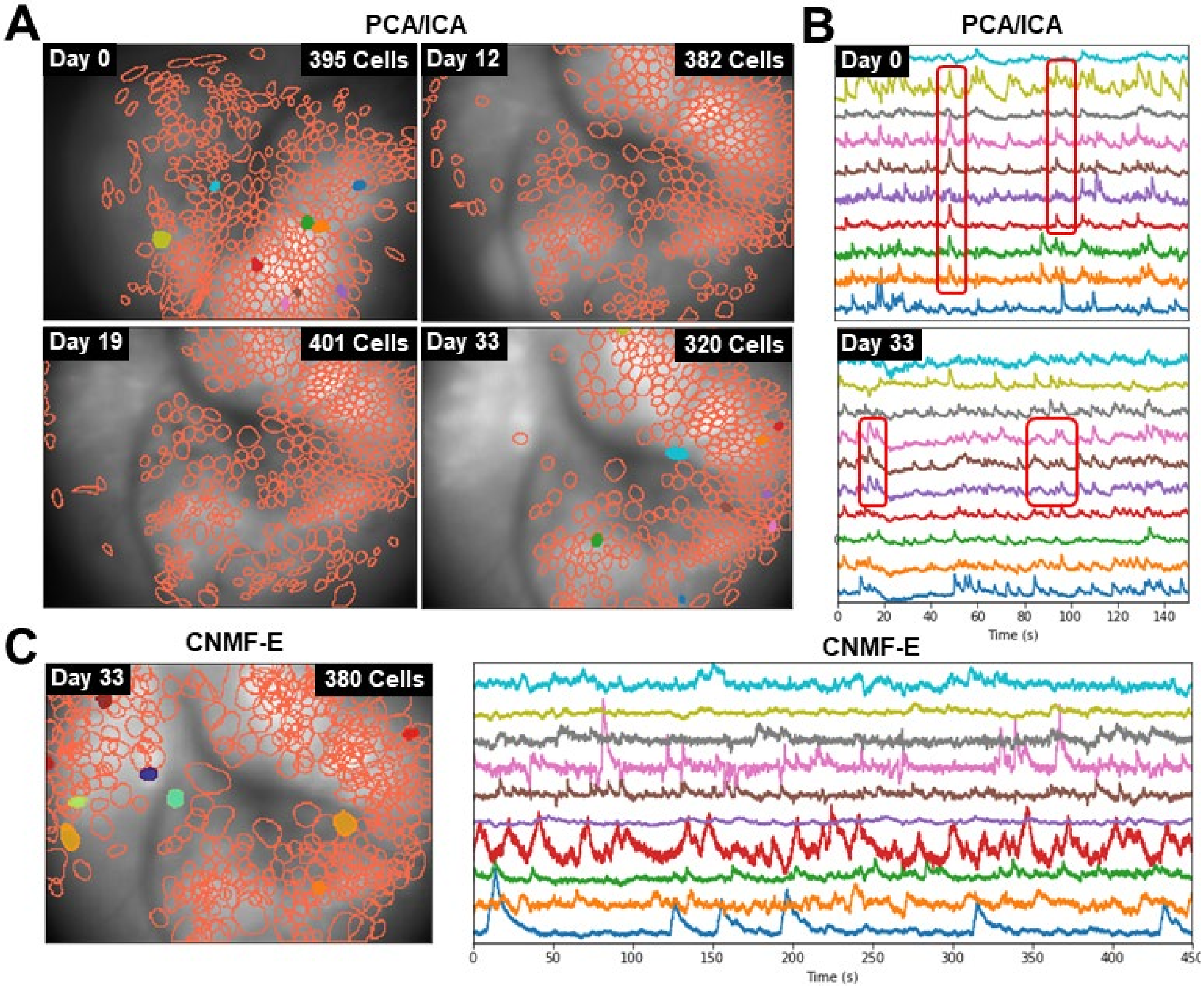


**Figure S4. Difference in extracted cell activity between PCA/ICA and CNMF-E performed on calcium imaging data obtained in an open-field arena. A)** Representative mouse showing that the number of striatal neurons identified by PCA/ICA was consistent over the course of the study as well as the failure of PCA/ICA to separate overlapping cell bodies. **B)** Neural signals extracted from 10 cells showing highly correlated activity (outlined in red) between nearby cells. **C)** CNMF-E was able to separate overlapping cells and did not exhibit highly correlated activity between cells.

***Comparison of Neural Signal Identification Techniques***

Calcium imaging data were loaded into Inscopix Data Processing Software (IDPS) (Inscopix, Palo Alto, CA) for preprocessing prior to identification of neural signals using region of interest (ROI) analysis, principal and independent component analysis (PCA/ICA)[58], or constrained non-negative matrix factorization for microendoscope data (CNMF-E)[59,60]. All data were first spatially down sampled by a factor of two, which is common practice in order to decrease processing time for subsequent analyses[55,56]. ROI analysis and PCA/ICA were performed using IDPS. ROI analysis was performed by manually selecting regions in the $\Delta F/F$ data that resembled cell body morphology and exhibited changes in relative fluorescence with respect to the surrounding background[61,62]. PCA/ICA was performed using both the default algorithm and parameters from IDPS and a spatiotemporal de-mixing value of 0.1[58]. In this analysis, the number of cells was visually estimated using the same criteria used for ROI analysis; the number of principal components was chosen to be 20% more than the number of cells identified[55,63–65]. We also evaluated the performance of ROI analysis and PCA/ICA after filtering the data with a gaussian spatial bandpass filter. The cut-off frequencies for this bandpass filter were independently selected for each data set based on the size of in-focus cell bodies. This led to cut-off frequencies ranging from 5 to 13 pixels for the high pass cut-off and 25 to 35 pixels for the low pass cut-off.

To perform analysis via CNMF-E, data were downsampled within IDPS by an additional factor of three for a total effective downsampling of 6x. Data were then exported as TIF files from IDPS and loaded into Matlab® (Natick, MA). To identify neurons via CNMF-E following the recommended procedure from Zhou et al. (2018)[59] and results were computed using Neuroscience Gateway[66]. Briefly, analysis was performed in patch mode using the largest possible patch sizes based on the available memory (128 GB) of the computational node. The ring model of background fluorescence[59] was used for all data sets with neuron and ring sizes selected based on the data set so that ring diameter was approximately 1.5 times the largest neuron diameters and described n Zhou et al. (2018)[59].

To compare the performance of different methods for isolating calcium activity given the presence of stimulation induced fluorescence from out of focus neurons, we overlaid the simulated effect of 30 out of focus neurons over real calcium imaging data obtained under anesthesia. The fluorescence of these neurons was spatially distributed according to symmetric 2D gaussians with a standard deviation of 50 pixels centered at random locations within the field of view. The 50-pixel standard deviation was chosen so that the diameter of an out-of-focus neurons was approximately 4 times the diameter of an in-focus neuron. The fluorescence intensity of each neuron was independently simulated according to Vogelstein et al. (2009)[68]. The activity rate of the simulated traces increased during stimulation (Figure S1). Simulated background fluorescence due to out-of-focus neurons was added to real calcium imaging data obtained under anesthesia. Simulated data were used to determine the effectiveness of each analysis technique for identifying cells in the presence of changes in out-of-focus fluorescence during electrical stimulation. ROI analysis, PCA/ICA and CNMF-E were then repeated on data with added simulated background fluorescence contamination and the results compared to uncontaminated data.

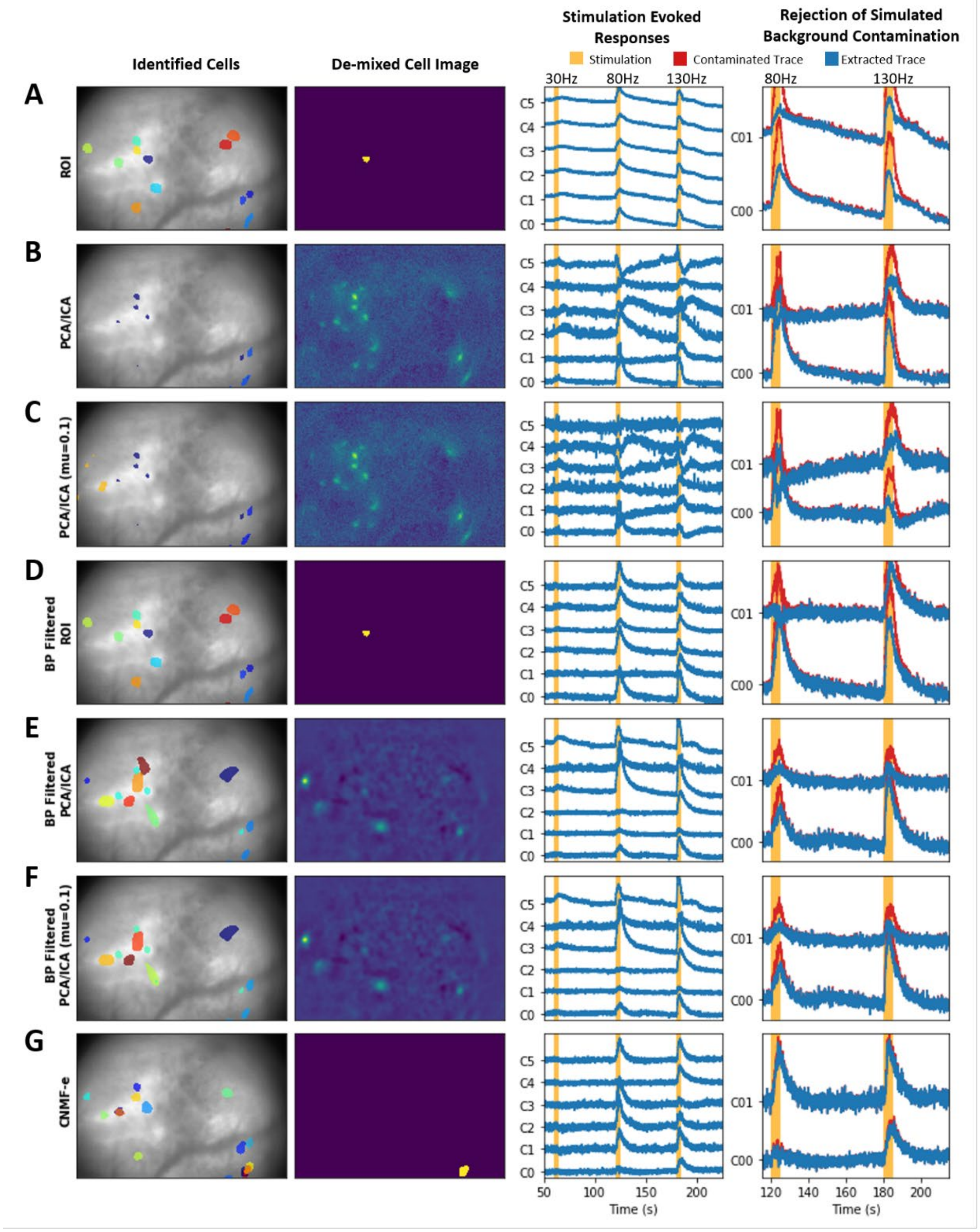


**Figure S5. Additional data extraction techniques. A)** The first column shows individual cells identified by ROI analysis. The second column shows a representative cell image obtained by ROI analysis showing a single well-defined cell body that was manually selected. The third column shows calcium traces during 30, 80, and 130 Hz stimulation trains that were extracted using region of interest (ROI) anaylsis. Lastly, the fourth column shows calcium traces extracted using ROI analysis during two stimulations, plotted on top of extracted traces from the same dataset and extracted using the same data analysis technique but contaminated with simulated background flourescence data. The same analysis is shown for other techniques including **B)** principle component analsysis followed by independent component analysis (PCA/ICA), **C)** PCA/ICA with spatiotemporal de-mixing value (mu) of 0.1, **D)** ROI analyis performed following band-pass filtering of the data, **E**) PCA/ICA following band-pass filtering, **F)** PCA/ICA (mu=0.1) following band-pass filtering, and **G)** constrained non-negative matrix factorization for microendoscope data (CNMF-E). These analysis were used to qualitatively assess the capability of each data analysis technique to identify spatially compact neurons, using the identified cell-bodies and de-mixed cell images, and to reject simulated background contamination
